# The Clinical Response of Upadacitinib and Risankizumab is Associated with Reduced Inflammatory Bowel Disease Anti-TNFα Inadequate Response Mechanisms

**DOI:** 10.1101/2022.05.16.492167

**Authors:** Jing Wang, Michael Macoritto, Heath Guay, Justin W. Davis, Marc C. Levesque, Xiaohong Cao

## Abstract

**Background and Aims:** JAK1 inhibitor upadacitinib and IL23 inhibitor risankizumab are efficacious in inflammatory bowel disease (IBD) patients who are anti-TNFα inadequate responders (TNF-IR). We aimed to understand the mechanisms mediating the response of upadacitinib and risankizumab.

**Methods:** Eight tissue transcriptomic datasets from IBD patients treated with anti-TNFα therapies along with single-cell RNAseq data from ulcerative colitis were integrated to identify TNF-IR mechanisms. RNAseq colon tissue data from clinical studies of TNF-IR Crohn’s disease patients treated with upadacitinib or risankizumab were used to identify TNF-IR mechanisms that were favorably modified by upadacitinib and risankizumab.

**Results:** We found seven TNF-IR up-regulated modules (M1-M7) related to innate/adaptive immune responses, interferon signaling and tissue remodeling, and five TNF-IR down-regulated modules (M8-M12) primarily related to metabolism. TNF-IR up-regulated cell types were inflammatory fibroblasts, post-capillary venules, inflammatory monocytes, macrophages, dendritic cells, and cycling B cells while subtypes of immature enterocytes, WNT5B+ cells and myofibroblasts were TNF-IR down-regulated cell types. Upadacitinib was associated with a significant decrease in the expression of most TNF-IR up-regulated modules in JAK1 responders (JAK1-R); in contrast, there was no change in these modules among TNF-IR patients treated with a placebo or among JAK1 inadequate responders (JAK1-IR). In addition, four of the six TNF-IR up-regulated cell types were significantly decreased after upadacitinib treatment in JAK1-R but not among subjects treated with a placebo or among JAK1-IR patients. We observed similar findings from colon biopsy samples from TNF-IR patients treated with risankizumab.

**Conclusions:** Collectively, these data suggest that upadacitinib and risankizumab affect TNF-IR up-regulated mechanisms, which may account for their clinical response among TNF-IR IBD patients.

## Introduction

Inflammatory bowel disease (IBD) consists primarily of ulcerative colitis (UC) and Crohn’s disease (CD), which are characterized by chronic relapsing inflammation in the gastrointestinal tract^1^. Although anti-tumor necrosis factor α (anti-TNFα) therapy was a breakthrough for IBD treatment, up to 40% of IBD patients are primary anti-TNFα treatment inadequate responders (TNF-IR) and another 30-40% lose response within one year of treatment^2, 3^. As such, there is a significant unmet need for new targeted therapies for IBD TNF-IR patients^4–10^.

The Janus kinase/signal transducers and activators of transcription (JAK/STAT) signaling pathway is implicated in the activation of inflammatory responses^11^ and many cytokines involved in IBD pathology transduce intracellular signals by this pathway. Several JAK inhibitors are currently approved or being evaluated for the treatment of IBD^3^ and showed clinical response for TNF-IR patients^12–14^. For example, the JAK1/3 inhibitor tofacitinib, the first oral JAK inhibitor approved for adult UC^15^, can induce and maintain clinical efficacy, endoscopic improvement and remission for TNF-IR patients based on results from the phase 3 OCTAVE Sustain study^14^. Upadacitinib is a selective inhibitor of JAK1, and showed efficacy in moderate-to-severe UC and CD TNF-IR patients based on the phase 2 U-ACHIEVE study^13^ and the phase 2 CELEST study, respectively^12^. Although Aguilar et al.^16^ generated an analysis of RNA sequencing (RNAseq) data from CD patients treated with upadacitinib in the phase 2 CELEST study, these investigators only used qPCR data from 19 cell markers to compare upadacitinib with anti-TNFα treatment. Thus, a more complete understanding of the mechanisms mediating JAK inhibitor response in TNF-IR IBD patients is missing.

Interleukin 23 (IL23) is a heterodimeric cytokine that can promote development of Th17 cells and cytokine-related immune responses and thus plays an essential role in the pathogenesis of IBD^17^. The anti-IL12/23 monoclonal antibody ustekinumab was approved for the treatment of moderate-to-severe CD and UC patients resistant to TNF antagonists^18, 19^. The IL23 inhibitors, risankizumab and brazikumab, also showed high response rates in anti-TNFα resistant CD patients ^20, 21^. Visvanathan et al.^22^ generated the first risankizumab transcriptomic dataset based on an analysis of a phase II CD clinical trial (ClinicalTrials.gov number NCT02031276). Although Visvanathan et al.^22^ found that risankizumab inhibited TNF expression compared with placebo, there was no significant difference in TNF expression between anti-IL23 responders (IL23-R) and anti-IL23 inadequate responders (IL23-IR) at week 0 and week 12. Peters et al.^23^ generated a ustekinumab transcriptomic dataset (GSE100833) from a phase 2b CD clinical trial of 87 anti-TNFα refractory patients. However, this data only included analysis of samples at week 0 prior to drug treatment. Therefore, no anti-IL23 study has yet identified molecular and cellular mechanisms mediating the clinical response of IL23 inhibitors in TNF-IR IBD patients.

Therefore, we integrated eight anti-TNFα treatment IBD transcriptomic datasets and a UC single-cell RNAseq dataset to identify the cellular and molecular mechanisms associated with an inadequate response to anti-TNFα therapies. We then used longitudinal upadacitinib RNAseq data from Aguilar et al.^16^ and risankizumab RNAseq data from Visvanathan et al.^22^ to identify TNF-IR mechanisms that were favorably modified by upadacitinib and risankizumab treatment and treatment.

## Materials and Methods

A schematic diagram of the study design is shown in Figure 1. Eight IBD transcriptomic datasets (100 TNF-R and 88 TNF-IR patients prior to treatment) (see Table 1) and one UC single-cell RNAseq dataset were integrated to reveal the consensus cellular and molecular mechanisms underlying sustained inflammation in TNF-IR IBD patients. Three of eight transcriptomic datasets also included control and post-treatment samples, which were used to validate the anti-TNFα mechanisms. RNAseq data from upadacitinib and risankizumab phase II CD clinical trials were used to support the clinical response of upadacitinib and risankizumab in IBD TNF-IR patients based on anti-TNFα mechanisms.

**Figure 1.**
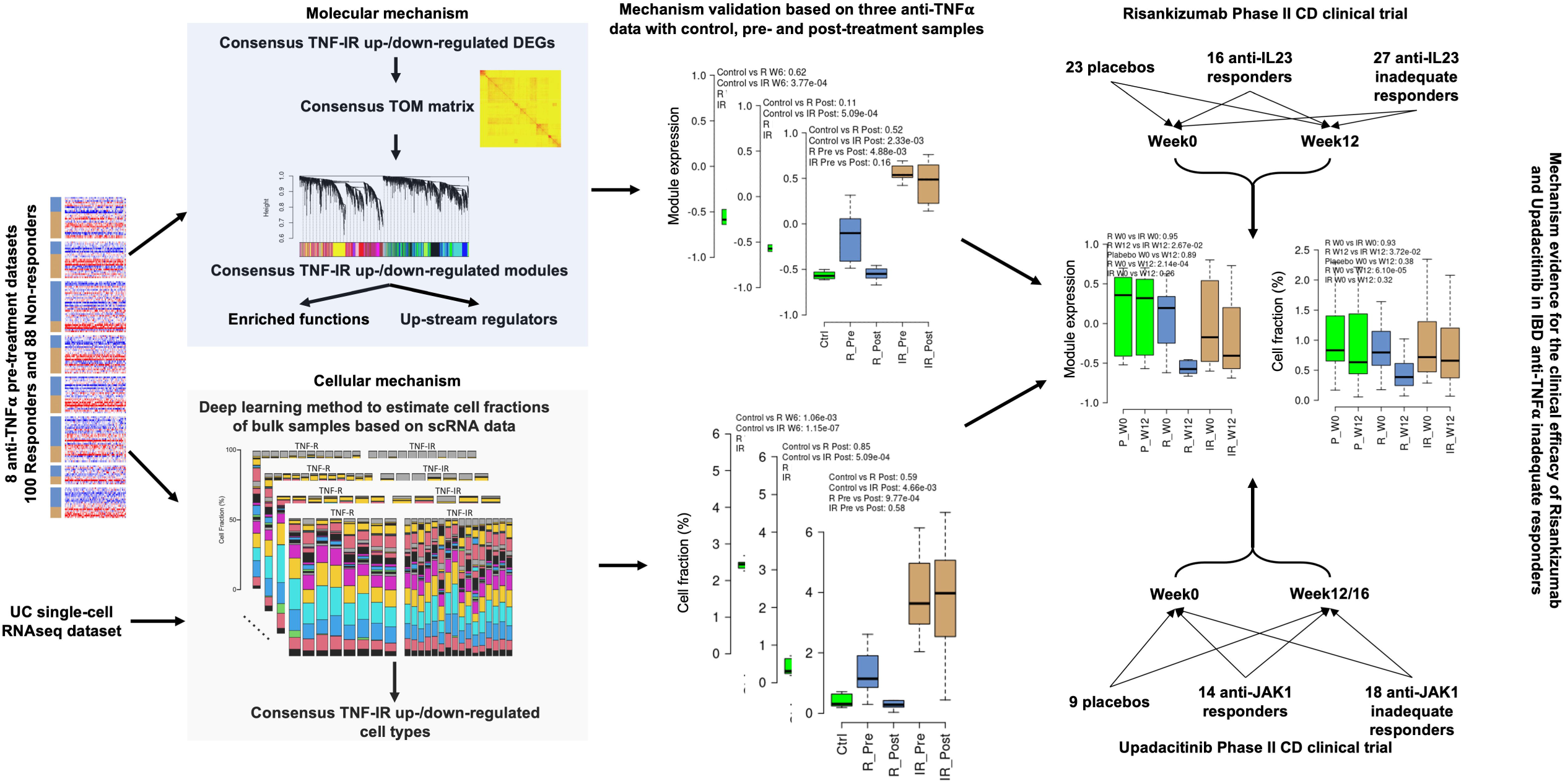
Overview of the integrative transcriptomics approach used for our analysis. We used eight anti-TNFα treatment data sets (100 responders and 88 non-responders) and a consensus network analysis and deep learning-based method to identify molecular and cellular mechanisms of anti-TNFα non-response that were then validated using independent control and post-treatment samples from three anti-TNFα treatment studies. RNAseq data from upadacitinib and risankizumab phase II CD clinical trials were used to validate the relationships and support the clinical response of upadacitinib and risankizumab in IBD TNF-IR patients. TNF-IR: TNF inadequate response; DEG: Differentially Expressed Gene; TOM: Topological Overlap Matrix.

**Table 1:**
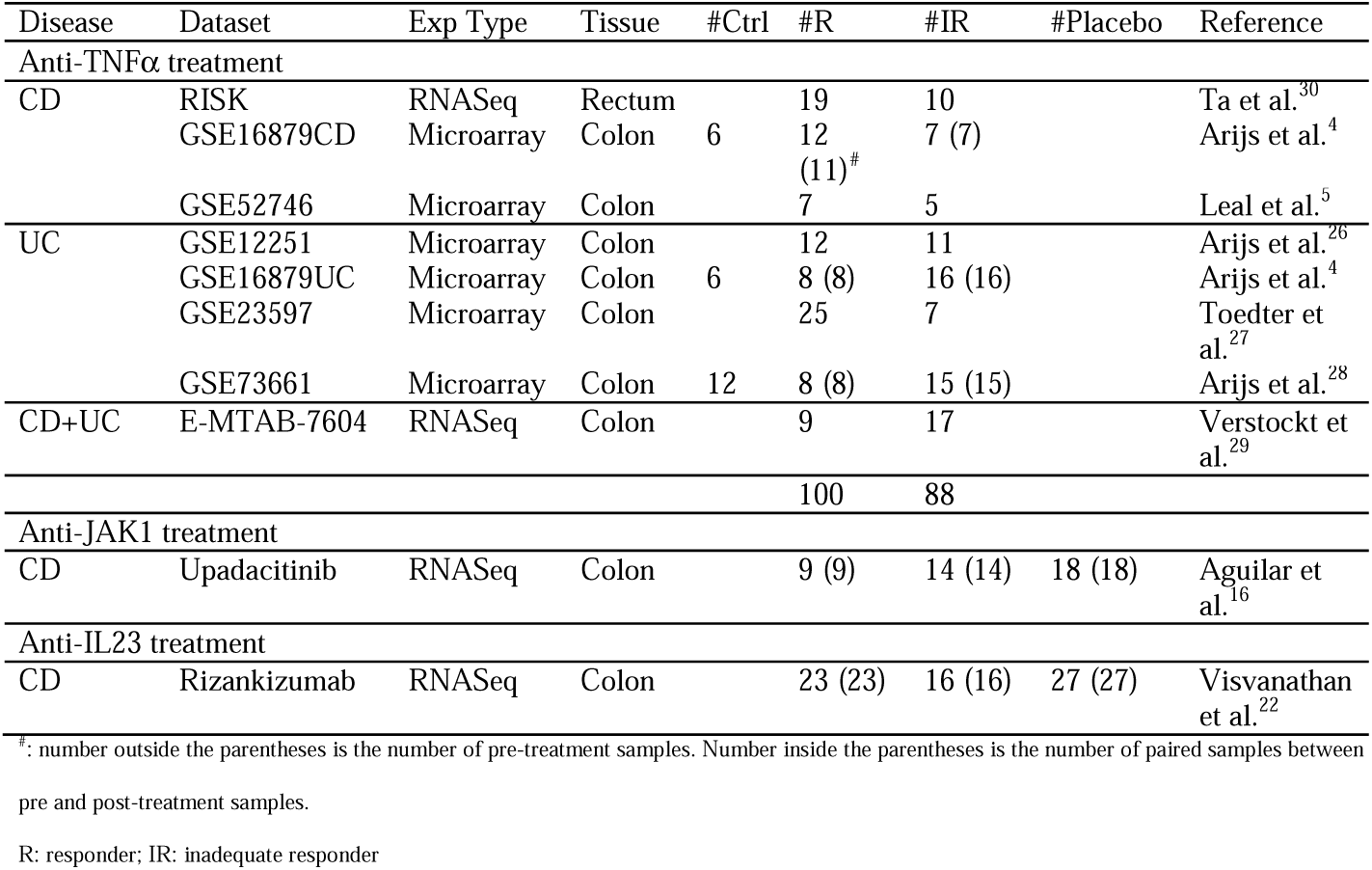
Bulk transcriptomic datasets used in this study.

### Anti-TNFα Gene Expression Data Sets

We searched Gene Expression Omnibus (GEO)^24^ and ArrayExpress^25^ for studies of IBD patients treated with anti-TNFα therapies that included at least five pre-treatment biopsies in each of the TNF-R and TNF-IR groups. We selected GSE16879 (CD only)^4^, GSE16879 (UC only), GSE52746^5^, GSE12251^26^, GSE23597^27^, GSE73661^28^ and E-MTAB-7604^29^ (see Table 1). GSE16879 CD, GSE16879 UC and GSE73661 also included biopsies from controls and paired post-treatment biopsies. We also included RISK^30^ data from the IBD Plexus program. Because RNA expression patterns of colon and ileum biopsies were significantly different (see Supplementary Figure 1), we only analyzed colon biopsies from GSE16879CD, RISK and E-MTAB-7604. The methods used for data processing can be found in the Supplementary Methods. Gene expression in inflamed mucosa from CD and UC were remarkably similar^31^. Thus, CD and UC data sets were considered together for meta-analyses.

### Consensus Differentially Expressed Genes (DEGs) Identification

Based on 21,655 genes included in at least five datasets, DEGs between TNF-IR subjects and TNF-R subjects were identified from each of the eight datasets of anti-TNFα treated IBD patients using the R package Limma^32^. Next, TNF-IR up-regulated (or down-regulated) meta-p-values and meta-FDRs were calculated based on the R packages metap and p.adjust (see Supplementary Methods). Finally, consensus DEGs were identified based on two rules: TNF-IR up-regulated (or down-regulated) meta-FDR<0.05 and one-sided p-value<0.2 in at least 5 data sets.

### Consensus WGCNA Modules Identification

Based on the consensus DEGs, the consensus WGCNA (weighted gene co-expression network analysis) method developed by Langfelder et al.^33^ was used to identify consensus gene modules across the eight anti-TNFα treated IBD patient datasets (see Supplementary Methods). Module expression was computed using the GSVA method^34^. A p-value comparing module expression and treatment status in each dataset was calculated by Wilcoxon rank-sum test^35^. A meta-p-value and meta-FDR were calculated in the same manner described for consensus DEG identification. TNF-IR up-regulated (or down-regulated) modules were identified using TNF-IR up-regulated (or down-regulated) meta-FDR<0.05 and one-sided p-value<0.2 in at least 5 datasets. Wilcoxon rank-sum paired tests were used to compare module expression between pre- and post-treatment samples. R package WebGestaltR^36, 37^ was used to identify enriched Gene Ontology terms for consensus modules with FDR<0.05 and the number of overlaps between module genes and genes in the GO term>5. Up-stream regulators of each module were predicted by IPA (Ingenuity Pathway Analysis)^38^ under FDR<0.05.

### Deep-Learning Based Cell Fraction Estimation Using Single-Cell RNAseq Data

The Scaden (single cell-assisted deconvolutional deep neural network (DNN)) method developed by Menden et al.^39^ was used to deconvolute the eight anti-TNFα IBD treatment bulk datasets based on 51 cell types from UC single-cell RNAseq data generated by Smillie et al.^10^ (see Supplementary Methods). Briefly, using 50,000 pseudo-bulk samples generated from UC single-cell RNAseq data, we built the 4-layer DNN model with L1 as the loss of function and Rectified Linear Unit (ReLU) activation for all layers except the last layer and we used softmax activation for the last layer (Supplementary Figure 2A). The model was trained and validated based on the leave-one-out subject method suggested by Menden et al.^39^. MAE (mean absolute error) and RSME (Root Mean Square Error) of the validation cohort was less than 0.012 and 0.024 (Supplementary Figure 2B and 2C). The cell fractions of 51 cell types were then estimated in each sample of the eight anti-TNFα IBD treatment bulk datasets based on the trained model. TNF-IR up-regulated (or down-regulated) cell types were identified based on the same meta-analysis used in the consensus WGCNA module identification. The mean of cell fractions in both TNF-R and TNF-IR groups for the significant cell types had to be over 0.5% in at least five data sets. Wilcoxon rank-sum paired tests were used to compare the cell fractions between pre- and post-treatment samples.

### Mechanism Validation Using the Data with Control, Pre- and Post-treatment Samples

We used three anti-TNFα datasets (GSE16879CD, GSE16879UC and GSE73661) with control, pre- and post-treatment samples to validate anti-TNFα mechanisms. The criteria for mechanism validation was 1) post-treatment module expression (or cell fractions) of TNF-R patients were not different (p>0.05) compared to controls while those of TNF-IR patients were significantly different compared to controls after treatment (p<0.05) or 2) if the post-treatment module expression (or cell fractions) of TNF-R patients were significantly different compared to controls, the module expression (or cell fractions) of TNF-R patients had to be significantly changed after treatment (p<0.05), while there was no change for TNF-IR patients after treatment (p>0.05). A module or cell type was considered a true anti-TNFα mechanism if it could be validated by at least two datasets based on the above criteria.

### Anti-TNF Mechanism Network Construction

To identify the relationship between anti-TNFα molecular and cellular mechanisms, genes in the TNF-IR up-regulated (or down-regulated) modules were compared with signatures of TNF-IR up-regulated (or down-regulated) cell types from Smillie et al.^10^. The following hypergeometric test^40^ was used to calculate the significance:

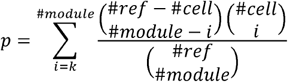

where *k* was the number of overlap genes between module and cell type, #module was the number of genes in the module, #cell was the number of genes in the cell signature and #ref was the union of genes in all modules and cell types. The significant relationships were identified as those with p<0.05 and *k*≥5.

### Longitudinal upadacitinib and risankizumab RNAseq Datasets

The longitudinal upadacitinib CD RNAseq dataset used in this study was from Aguilar et al.^16^ and we analyzed the paired week 0 and week 12/16 samples from TNF-IR patients and colonic biopsies. JAK1-R and JAK1-IR patients were defined based on clinical response at week 12/16 (≥30% reduction from week 0 in stool frequency and/or abdominal pain score and both not greater than week 0 score)^12^. Because the 3mg BID arm did not have any JAK1-R patients, we removed this arm in our analysis (Supplementary Figure 3A). We combined the 6mg BID, 12mg BID, 24mg BID and 24mg QD arms to increase statistical power. This resulted in paired week 0 and week 12/16 samples from 9 placebo subjects, 14 JAK1-R patients and 18 JAK1-IR patients.

The longitudinal risankizumab CD RNAseq dataset used in this study was from Peters et al.^23^ and we selected the paired week 0 and week 12 samples from TNF-IR patients and colonic biopsies. IL23-R and IL23-IR patients were defined based on the Crohn’s Disease Endoscopic Index of Severity (CDEIS)^41^. We combined the 200mg IV and 600mg IV arms during analyses (Supplementary Figure 3B) and this resulted in paired week 0 and week 12 samples from 23 placebo subjects, 16 risankizumab-treated IL23-R patients and 27 IL23-IR patients. upadacitinib and risankizumab RNAseq datasets were processed based on the same method as E-MTAB-7604 and RISK.

## Results

### Molecular Mechanisms of Anti-TNFα Inadequate Response

Based on meta-analyses from eight anti-TNFα pre-treatment datasets, 1,972 consensus TNF-IR up-regulated genes and 2,021 TNF-IR down-regulated genes were identified under meta-FDR<0.05 and individual dataset one-sided p<0.2 in at least 5 datasets (Supplementary Table 1-2). Twelve consensus network modules (M1-M12, Supplementary Table 3) were identified based on the consensus genes. Most of the genes in each module had consistent expression patterns across both CD and UC datasets (Figure 2A), which suggested that CD and UC had similar TNF-IR mechanisms. We correlated M1-M12 module expression with treatment status in each dataset and seven modules were identified as consensus TNF-IR up-regulated modules and five modules were consensus TNF-IR down-regulated modules (Figure 2B, Supplementary Table 4-5). Figure 2C shows the representative Gene Ontology (GO) terms enriched in each module (all enriched GO terms can be found in Supplementary Tables 6-17).

**Figure 2.**
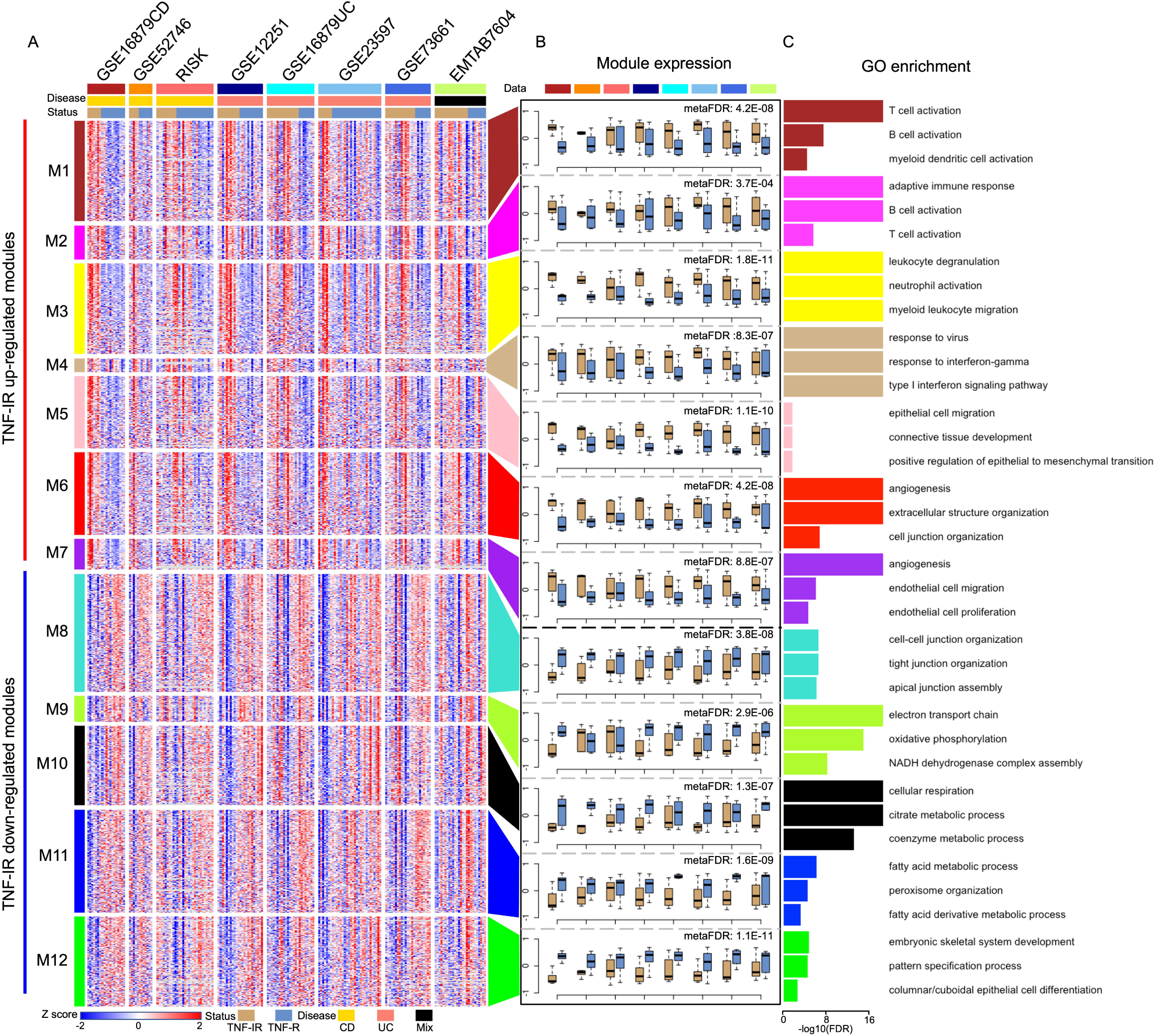
Identification of TNF-IR up- and down-regulated modules. (A) Heatmap of z-transformed expression of consensus DEGs in eight data sets. Genes are clustered based on 12 anti-TNFα related modules. (B) Module expression in eight data sets. Blue and light brown boxes represent TNF-R and TNF-IR patients, respectively. Meta-FDR is calculated based on the comparison of module expression between TNF-IR and TNF-R patients in each data set. The order of the data sets in (B) is the same as that in (A). (C) Functional annotation of genes in each module based on Gene Ontology enrichment. Three representative enriched processes are shown, sorted by p-value.

TNF-IR up-regulated module M1 was highly related to adaptive and innate immune responses including B, T and myeloid dendritic cell activation. Module M2 was mainly related to adaptive immune responses including B and T cell activation. Module M3 was associated with innate immune responses with genes related to myeloid leukocyte migration and neutrophil activation. Module M4 was associated with interferon signaling including response to interferonγ and type I interferon signaling. Modules M5-M7 were associated with tissue remodeling; module M5 was related to connective tissue development, epithelial cell migration and positive regulation of epithelial to mesenchymal transition; module M6 was related to angiogenesis, extracellular structure organization and cell junction organization and module M7 was related to angiogenesis, cell junction assembly and epithelial cell migration.

Modules M8-M12 were consensus TNF-IR down-regulated modules. Module M8 was associated with barrier function including cell-cell junction organization, tight junction organization and apical junction assembly. Modules M9-M11 were related to metabolism; module M9 was associated with energy metabolism (electron transport chain, oxidative phosphorylation and NADH dehydrogenase complex assembly); module M10 was associated with cellular respiration, citrate and coenzyme metabolic processes and module M11 was associated with lipid metabolism (fatty acid metabolic processes, peroxisome organization, fatty acid derivative metabolic processes). Module M12 was characterized by developmental functions including embryonic skeletal system development, pattern specification processes and columnar/cuboidal epithelial cell differentiation.

Upstream regulator analysis identified 218 regulators related to at least two TNF-IR up-regulated modules (Supplementary Table 18) that included several well-known TNF-IR associated genes (e.g. OSM^42^, TREM1^8^, IL1B^43^). Relationship between TNF-IR up-regulated modules and regulators can be also found in Figure 4B. We also found 31 regulators related to at least two TNF-IR down-regulated modules (Supplementary Table 19). **Anti-TNF**α **Treatment was Associated with Reduced Molecular Mechanisms in Anti-TNF**α **Responders** GSE16879CD, GSE16879UC and GSE73661 also included control and post-treatment samples (Table 1) and were used as independent datasets to validate the different effects of anti-TNFα treatment on molecular mechanisms in TNF-R and TNF-IR patients. As shown in Figure 3A, TNF-IR up-regulated modules M1, M2, M4, M6 and M7 were validated in GSE16879CD based on Criterion One that post-treatment expression of modules in TNF-R patients, but not TNF-IR patients, were similar to those in controls (p>0.05). In GSE16879CD, modules M3 and M5 were validated based on Criterion Two that expression of modules in TNF-R patients, but not TNF-IR patients, were significantly decreased after anti-TNFα treatment (p<0.05), even if not reduced to the levels of control subjects. In addition, five TNF-IR up-regulated modules in GSE16879UC (except modules M3 and M7) and all 7 in GSE73661 were validated based on Criterion Two (Figures 3B & 3C). For the five TNF-IR down-regulated modules, supplementary Figure 4 showed that all these modules were validated in at least two datasets based on Criteria One or Two.

**Figure 3.**
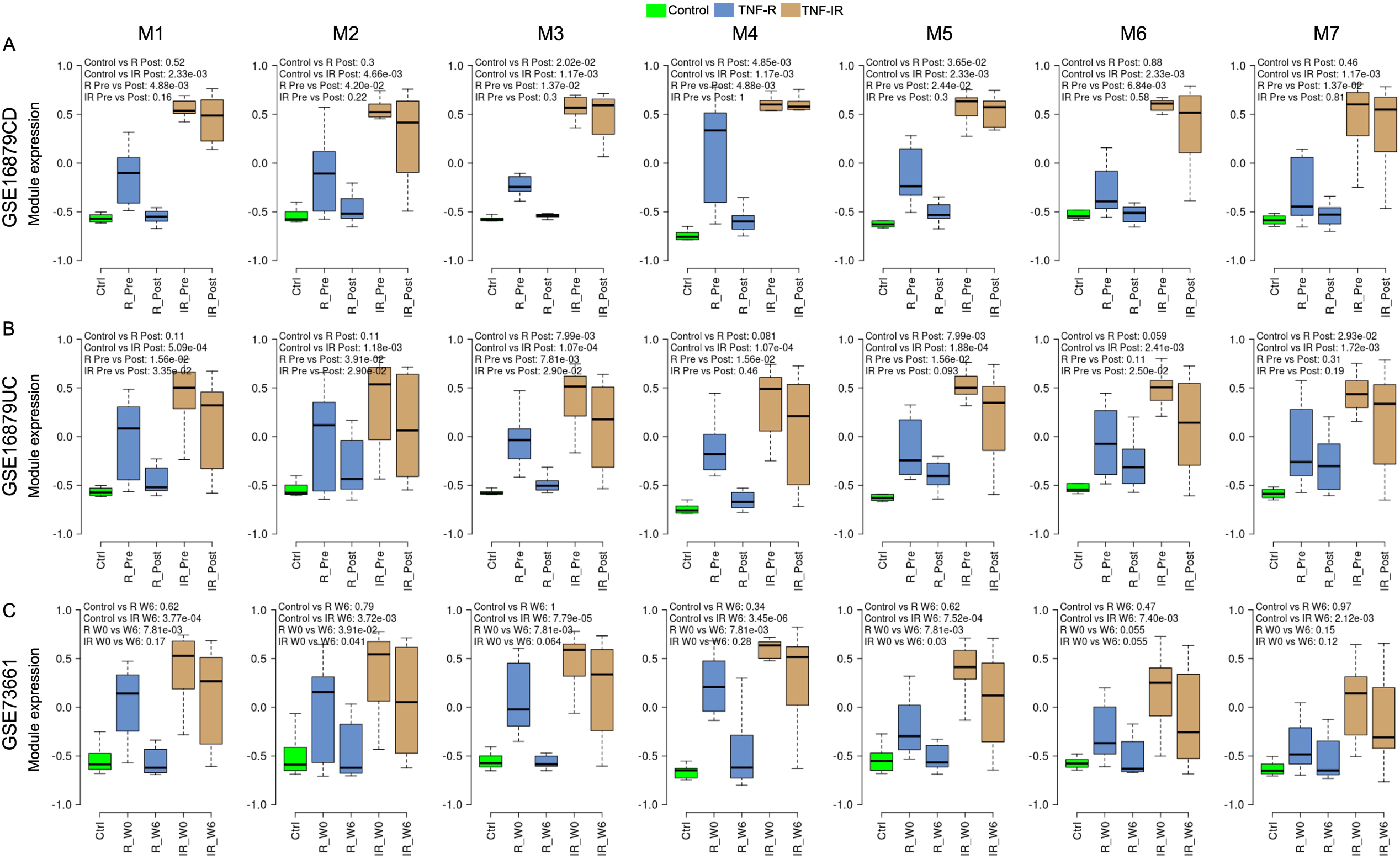
The expression of TNF-IR up-regulated modules could not be returned to normal levels after anti-TNFα treatment in non-responders based on the GSE16879CD (A), GSE16879UC (B) and GSE73661 (C) datasets. Green, blue and light brown boxes in the boxplots represent controls, TNF-R and TNF-IR patients, respectively. The p-values in the top-left of boxplots were calculated based on Wilcoxon rank sum test.

**Figure 4.**
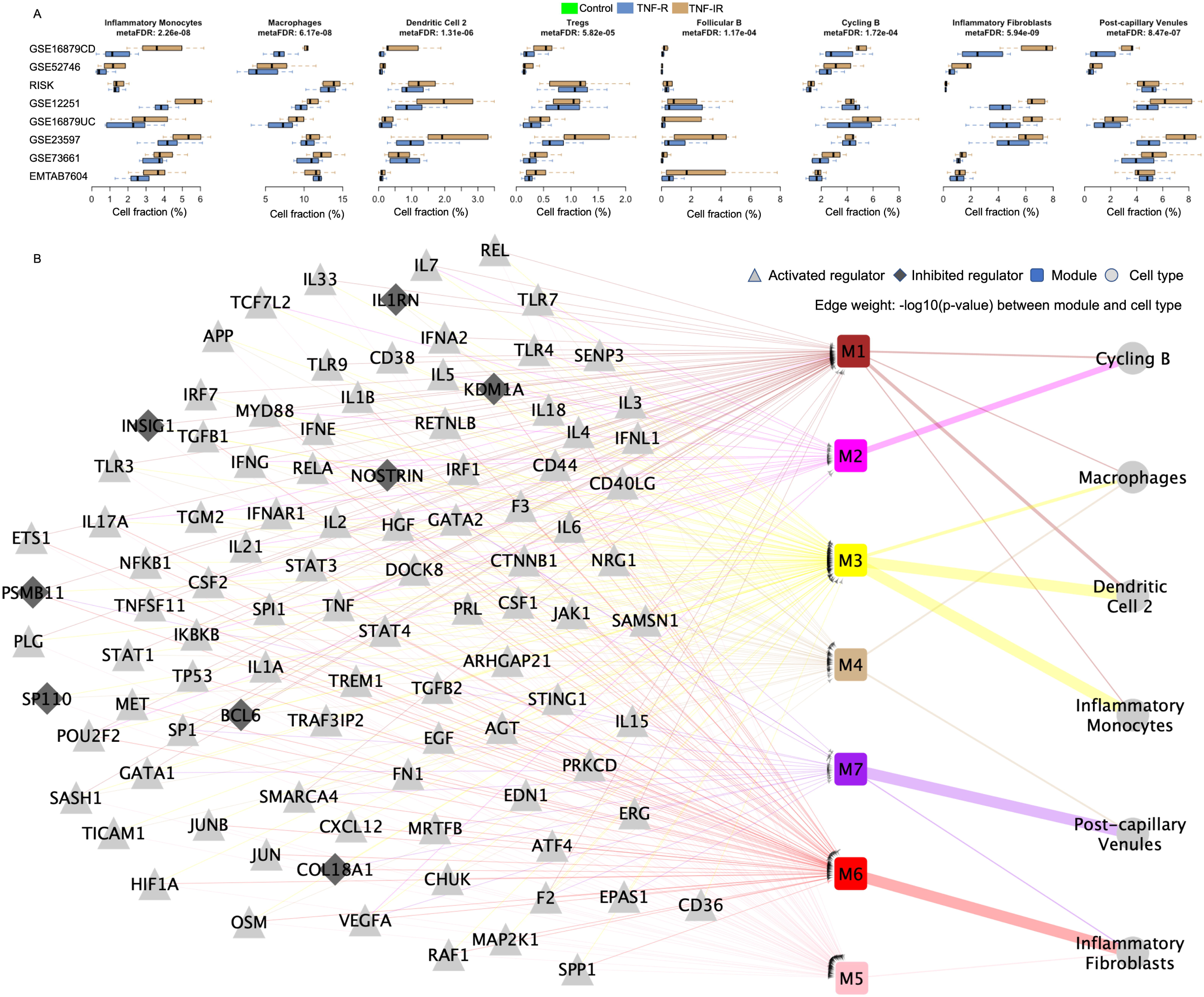
TNF-IR up-regulated cell type identification (A) and relationship among TNF-IR up-regulated modules, cell types and upstream regulators (B). Meta-FDR in (A) was calculated based on the comparison of cell fractions between TNF-IR and TNF-R patients in eight data sets. Triangle, diamond, square and circle in (B) represented activated regulator, inhibited regulator, module and cell type, respectively. Regulators related to at least 3 modules were visualized in (B). Module colors in (B) were the same with those in Figure 2A. Edge weight in (B) represented −log10(p-value) between module and cell type. Edge colors were same with the colors of the linked modules.

### Cellular Mechanisms of Anti-TNFα Inadequate Response

Cell fractions of 51 cell types in eight anti-TNFα treatment datasets were estimated based on the deep-learning model created by Menden et al.^39^ using UC single-cell RNAseq data^10^. As shown in Figure 4A, eight cell types, as defined by Smillie et al.^10^ (inflammatory fibroblasts, inflammatory monocytes, macrophages, post-capillary venules, dendritic cells 2, Tregs, follicular and cycling B cells) were identified as TNF-IR up-regulated cell types and had significantly higher cell fractions in TNF-IR patients compared to TNF-R patients in most of the eight datasets. We identified six TNF-IR down-regulated cell types (see Supplementary Figure 6A) that included four epithelial cell types (cycling TA, immature enterocytes 2, immature goblet and TA 2) and two fibroblast cell types (WNT5B+ 2 and myofibroblasts), as defined by Smillie et al.^10^.

Based on the three datasets we used for molecular mechanism validation (GSE16879CD, GSE16879UC, and GSE73661), we found that most TNF-IR up-regulated cell types could be validated using Criteria One or Two in at least two datasets (Supplementary Figure 5). Tregs and follicular B cells were the exceptions because the post-treatment fractions of Tregs in TNF-IR patients were similar to controls in all three datasets and follicular B cell fractions were not significantly changed after treatment for both TNF-R and TNF-IR patients. Among the six TNF-IR down-regulated cell types, immature enterocytes 2, WNT5B+ 2 and myofibroblasts were validated in at least two datasets (Supplementary Figure 6B). Therefore, in the following analyses, we only focused on cell types that were validated in at least two datasets (six TNF-IR up-regulated cell types and three TNF-IR down-regulated cell types).

### Relationship Between Anti-TNFα Molecular and Cellular Mechanisms

To cross-validate molecular and cellular mechanisms, we built anti-TNFα mechanism networks (Figure 4B). Module M1 was an adaptive and innate immune module that was highly related to all immune cell types. Module M2 was significantly related to cycling B cells and module M3 was associated with macrophages, inflammatory monocytes and dendritic cells 2. Modules M5, M6 and M7 were tissue remodeling modules highly related to inflammatory fibroblasts and post-capillary venules. Interferon signaling module M4 was related to macrophage and post-capillary venules and was identified by previous studies ^44, 45^. The TNF-IR down-regulated cell type immature enterocytes 2 was highly related to lipid metabolism module M11. There was no clear relationship between any of the TNF-IR down-regulated modules and two fibroblast cell types (WNT5B+ 2 or myofibroblasts).

### Upadacitinib and risankizumab Reduced TNF-IR Up-regulated Mechanisms among IBD Patients

We initially determined whether inhibiting JAK1 with upadacitinib or inhibiting IL23 with risankizumab impacted TNF mRNA abundance among IBD patients. As shown in Supplementary Figure 7, there was no significant difference in TNF mRNA abundance between week 0 and week 12/16 among placebo, JAK1-R, and JAK1-IR patients or between week 0 and week 12 among placebo and IL23-R patients (p>0.05). TNF mRNA abundance was significantly different between week 0 and week 12 for IL23-IR patients (p<0.05). These results indicated that the response of upadacitinib and risankizumab on TNF-IR patients was not directly related to TNF mRNA abundance.

Next, we determined whether upadacitinib or risankizumab had an effect on anti-TNFα mechanisms. As shown in Figure 5A, the expression of six TNF-IR up-regulated modules (except module M2) were significantly decreased in JAK1-R patients when comparing week 0 to week 12/16 (p<0.05). However, because of the slight decrease of module expression in JAK1-IR patients after upadacitinib treatment, no module was significantly different between JAK1-R and JAK1-IR patients at week 12/16. We observed similar results for the TNF-IR down-regulated modules (Supplementary Figure 8A). Cell deconvolution analyses using UC single-cell RNAseq data^10^ indicated that the cell fractions of inflammatory fibroblasts, inflammatory monocytes, macrophages, cycling B cells and immature enterocytes 2 were significantly affected after treatment in JAK1-R patients but no cell type was significantly different between JAK1-R and JAK1-IR patients at week 12/16 (Figure 5B and Supplementary Figure 8B).

**Figure 5.**
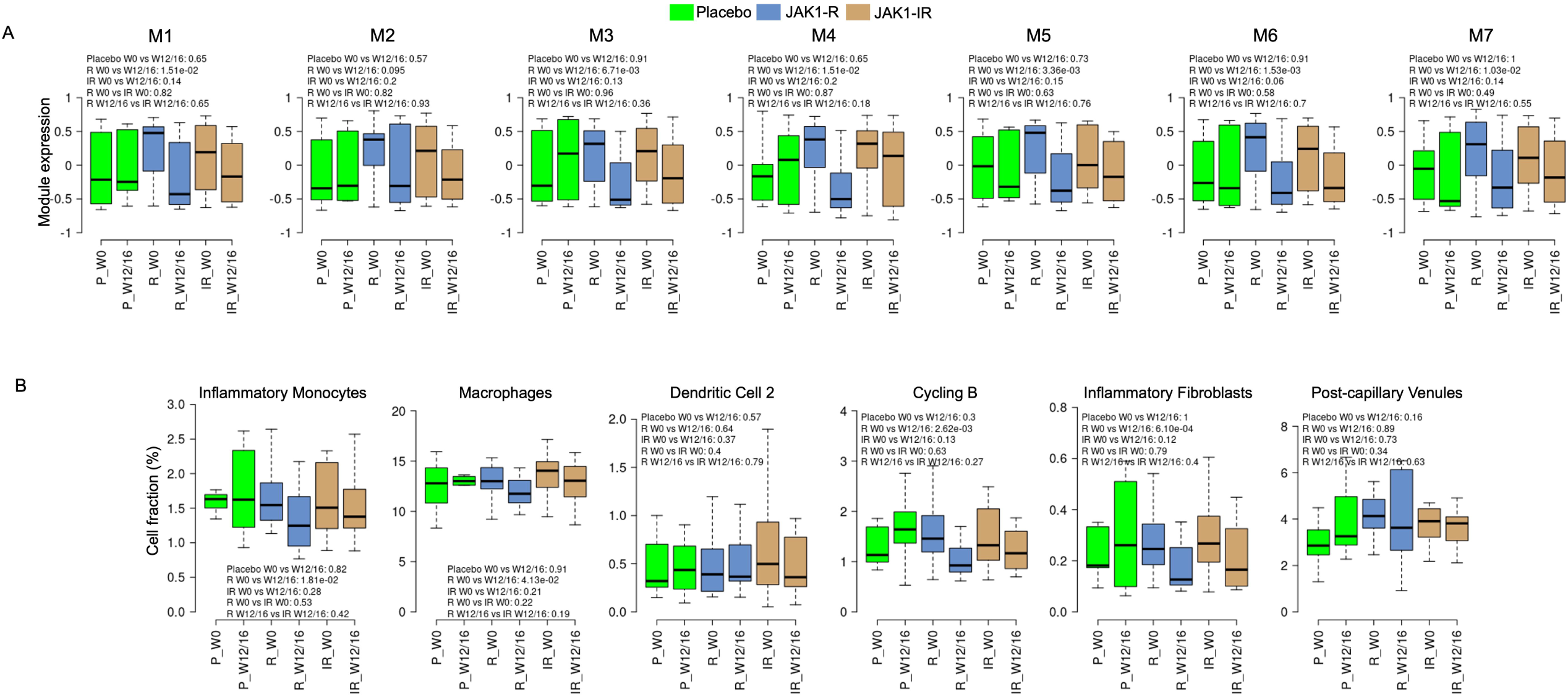
Upadacitinib significantly affected JAK1 related TNF-IR up-regulated modules and cell types in JAK1-R patients but not placebos and JAK1-NR patients. (A) Comparison of expression of seven TNF-IR up-regulated modules in placebos (green), JAK1-R (blue) and JAK1-NR (light brown) patients for week 0 and week 12/16. (B) Comparison of cell fractions of six TNF-IR up-regulated cell types in three groups for week 0 and week 12/16. The p-values were calculated based on Wilcoxon rank sum tests.

As shown in Figure 6A, after risankizumab treatment, the expression of all TNF-IR up-regulated modules were significantly decreased (p<0.05) in IL23-R patients. There was no significant change in TNF-IR up-regulated modules among patients treated with placebo or among IL23-IR patients, although the expression of modules M1-M5 were slightly decreased after risankizumab treatment. TNF-IR up-regulated modules M1 and M3-M5 were also significantly different (p<0.05) when comparing module expression between IL23-R and IL23-IR patients at week 12. The expression of all TNF-IR down-regulated modules was significantly increased (p<0.05) in IL23-R patients but there was no significant change in TNF-IR down-regulated module expression among patients treated with placebo and among IL23-IR patients (Supplementary Figure 9A). Expression of modules M8, M11 and M12, but not M9 and M10, were significantly different between IL23-R and IL23-IR patients at week 12 (Supplementary Figure 9A).

**Figure 6.**
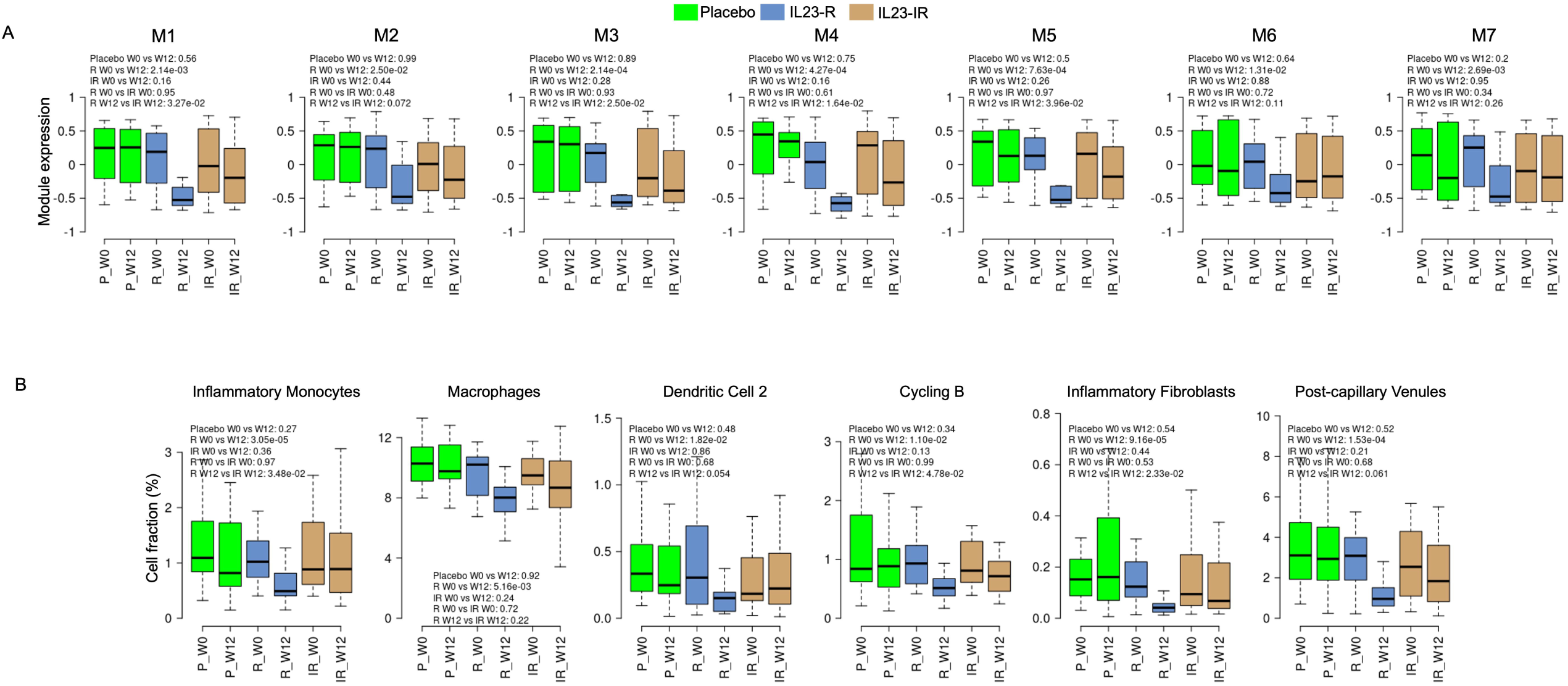
Risankizumab significantly affected IL23-related TNF-IR up-regulated modules and cell types in IL23-R patients but not placebos and IL23-IR patients. (A) Comparison of expression of seven TNF-IR up-regulated modules in placebo (green), JAK1-R (blue) and JAK1-NR (light brown) for week 0 and week 12. (B) Comparison of cell fractions of six TNF-IR up-regulated cell types in three groups for week 0 and week 12. The p-values were calculated based on Wilcoxon rank sum tests.

Cell deconvolution analyses indicated that six TNF-IR up-regulated cell types were not significantly different between IL23-R and IL23-IR patients at week 0, as expected, while these same six cell types were significantly decreased after treatment among IL23-R patients but not placebo or IL23-IR patients (Figure 6B). Inflammatory monocytes and inflammatory fibroblasts had significantly different cell fractions between IL23-R and IL23-IR patients at week 12. Among the three TNF-IR down-regulated cell types, only immature enterocytes 2 were significantly affected between weeks 0 and 12 by risankizumab in IL23-R patients. There was no significant difference between IL23-R and IL23-IR patients at week 12 (Supplementary Figure 9B).

## Discussion

The mechanisms associated with TNF inadequate response and the ability of upadacitinib and risankizumab to induce clinical improvement in TNF-IR IBD patients^12, 20^ are not fully understood. Integrating eight anti-TNFα treatment transcriptomic datasets, UC single-cell RNAseq dataset and two RNAseq datasets from upadacitinib and risankizumab clinical trials, we found a correlation between the clinical response to upadacitinib and risankizumab and effects on TNF-IR molecular and cellular mechanisms among anti-TNFα inadequate responders.

Several studies have tried to elucidate the molecular mechanisms responsible for sustained active inflammation in IBD despite anti-TNFα treatment^4, 5, 46^. However, to date only Liu et al.^46^ integrated multiple anti-TNFα treatment datasets to identify stable, reproducible TNF-IR mechanisms. Liu et al.^46^ integrated five datasets using ComBat methodology^47^ to reduce data specific batch effects and identified 274 TNF-IR up-regulated genes and 14 TNF-IR down-regulated genes. Comparing these genes with the genes in our 12 modules, we found that 149 and 111 of the up-regulated genes identified by Liu et al^46^ belonged to module M3 (innate immune response) and modules M5-M7 (tissue remodeling), respectively, while 12 of the 14 down-regulated genes identified by Liu et al^46^ belonged to module M12 (development functions) (Supplementary Table 20). None of the up-regulated genes identified by Liu et al^46^ were annotated by module M2 (adaptive immune response) which plays a major role in the pathogenesis of IBD^48^. A GO enrichment analysis of the up-regulated genes identified by Liu et al^46^ showed that most genes were related to the innate immune response and tissue remodeling pathways (Supplementary Table 21), and there was no significant GO term derived from the down-regulated genes identified by Liu et al^46^. These results suggest that the WGCNA pipeline developed in our study was more sensitive in identifying anti-TNFα IR mechanisms compared with the ComBat data integration method used by Liu et al^46^, which may be due to obscured differences between TNF-R and TNF-IR patients during removal of the batch effect.

Neutrophils are known to play an important role in the pathogenesis of IBD^49^. We found that TNF-IR up-regulated module M3 was related to neutrophil activation; anti-TNFα treatment significantly inhibited the expression of module M3 in TNF-R but not TNF-IR patients, which is consistent with a previous report^50^ and consistent with a recent publication detailing the role of neutrophils in IBD ulcerations in TNF-IR patients^43^. However, we cannot estimate the neutrophil cellular proportions in these samples due to the lack of a neutrophil cell cluster in the UC single-cell RNAseq data used in our study or in any other published IBD single-cell data^9, 51^. The lack of a neutrophil signature in our datasets may be due to the relatively low level of RNA content of neutrophils and the fact that these cells are prone to degradation using most RNA collection and purification methodologies^52^. The lack of a neutrophil signature is an area for development in the future as single-cell level data for neutrophils in IBD would greatly enhance our understanding of their role in disease.

In our study, the longitudinal RNAseq data from clinical trials of upadacitinib validated that JAK1 inhibition significantly decreased the expression and cell fractions of most TNF-IR up-regulated modules and cell types in JAK1-R patients while there was no change in placebo-treated patients and JAK1-IR patients; we observed similar findings for risankizumab. These results suggest that resolution of adaptive/innate immune signaling, and blockade of tissue remodeling may be important for disease resolution. When comparing between JAK-R and JAK-IR patients at week 12/16 or between IL23-R and IL23-IR patients at week 12, we found that four modules (M1, M3, M4, and M5) and three cell types (Inflammatory Monocytes, Cycling B and Inflammatory Fibroblasts) were significantly different for risankizumab but not upadacitinib.

Like our study, Aguilar et al.^16^ also identified cell types regulated by upadacitinib based on UC single-cell data^10^. Aguilar et al.^16^ compared the upadacitinib response signature to cell signatures and filtered the cell types based on the number of overlapping genes. As the cell signatures in the UC single-cell data are highly overlapping among similar subsets of cells (Supplementary Figure 10), the method used by Aguilar et al.^16^ was highly affected by the dependencies between closely related cell subtypes. For example, Aguilar et al.^16^ found that the median expression of cell signatures from each fibroblast cell type was reduced after upadacitinib treatment in JAK1-R patients. In contrast, the deep-learning based cell deconvolution method used in our study optimized features based on all genes in the single-cell data and avoided the problems associated with dependencies between closely related cell subtypes. In our findings, inflammatory fibroblasts were the only fibroblast cell type that was significantly reduced in JAK1-R patients after upadacitinib treatment (Supplementary Figure 11). Aguilar et al.^16^ found an inconsistent pattern among different fibroblast cell types with their qPCR results for subjects in the upadacitinib clinical study. Inflammatory fibroblast cell marker CHI3L1 was significantly reduced after treatment in JAK1-R patients while WNT5B+ cell markers PDGFD, PTGDR2 and SOX6 expression were slightly increased in JAK1-R patients. These results were consistent with our deep-learning cell deconvolution results (Supplementary Figure 11). The qPCR results from Aguilar et al.^16^ also validated the inconsistent pattern between inflammatory fibroblasts and WNT5B+ fibroblasts related to anti-TNFα treatment.

T cells are pivotal mediators of mucosal damage in both CD and UC^53^. However, among 11 T cell populations described by Smillie et al.^10^, the cell fractions of eight T cells were lower than 0.5% in both TNF-R and TNF-IR patients from at least five pre-anti-TNFα-treatment datasets (Supplementary Figure 12). Although CD8+ lamina propria (LP) and Cycling T cells had higher cell fractions, the comparisons between TNF-R and TNF-IR patients were not consistent across eight datasets. Thus, we only identified Tregs as a candidate up-regulated cell type among 11 T cell types. Because Tregs were unable to pass the validation criteria (Supplementary Figure 5), we did not include any T cells when integrating with the upadacitinib CD RNAseq data. We checked whether the cell fractions of T cells were affected by upadacitinib and found that cycling T cells were significantly decreased in JAK1-R patients but not among placebo-treated patients or JAK1-NR patients (Supplementary Figure 13), which was consistent with the qPCR results of Aguilar et al.^16^ that used IFNγ as a cell marker for this population of T cells.

One of the major limitations of our study is that we did not experimentally validate our findings related to TNF-inadequate responses and inadequate responses to upadacitinib and risankizumab. However, we validated our TNF-IR related findings based on the independent control and post-treatment samples from three anti-TNFα treatment datasets and we validated our clinical trial analyses using two different therapies (upadacitinib and risankizumab). Although blockade of some TNF-IR up-regulated mechanisms by upadacitinib and risankizumab are understandable mechanistically (such as blockade of OSM activation by JAK1 inhibition), other mechanisms affected by JAK1 or IL23 blockade are more difficult to explain. The results of the clinical studies also leave unanswered questions about how JAK1 inhibition and IL23 blockade result in relatively similar patterns of module and cell changes in TNF-IR patients. Another potential limitation of our study relates to the best ways to integrate results from single-cell-based methods and signature-based methods. Other methods for cell deconvolution have been developed and can estimate the proportions for more cell types than we examined in the current study. For example, signature-based cell deconvolution methods such as XCell^54^ can infer the abundance of 64 cell types (including neutrophils) based on 489 cell signatures. However, most of the signatures used by methods such as XCell are generated from peripheral blood datasets (e.g. Blueprint^55^), and therefore, methods like XCell, may be insufficient to estimate the cellular composition of tissue samples^56^. In summary, the upadacitinib and risankizumab CD clinical trial data suggests that upadacitinib and risankizumab may block pathways that remain active in IBD patients not responding to anti-TNFα therapy. This may mechanistically account for the response of JAK1 inhibitors and IL-23 targeted therapies among this difficult to treat population of patients. The mechanism identification pipeline we developed can also be used in other diseases to generate new hypotheses for other new treatments.

## Acknowledgements

The results published here are in part based on data obtained from the IBD Plexus program of the Crohn’s & Colitis Foundation.

