## Supplementary Figures for "The Clinical Response of Upadacitinib and Risankizumab is Associated with Reduced Inflammatory Bowel Disease Anti-TNFα Inadequate Response Mechanisms"

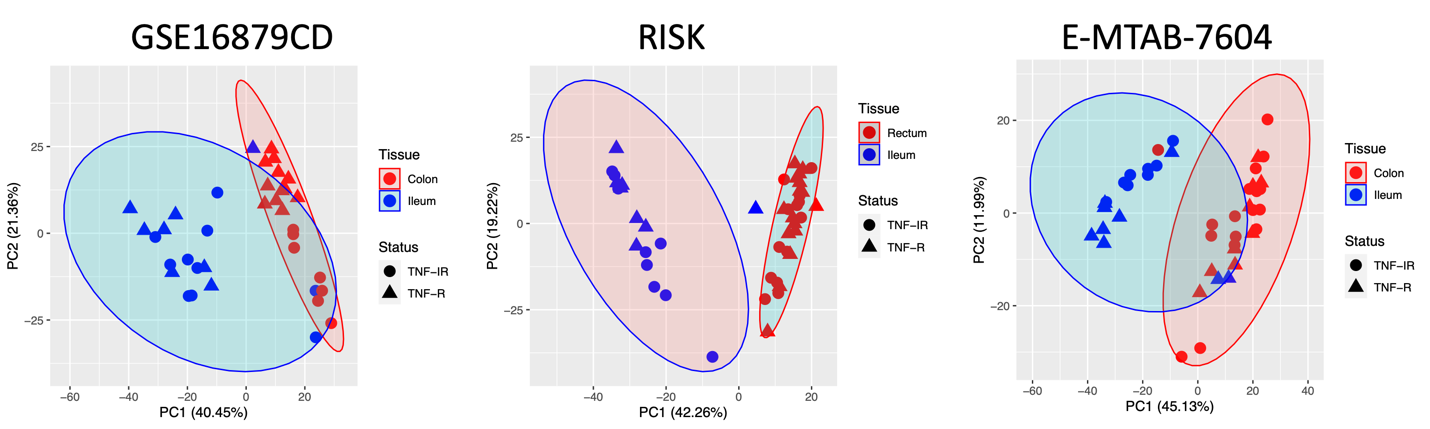

Supplementary Figure 1. PCA plots for colon/rectum and ileum samples in GSE16879CD, RISK and E-MTAB-7604 datasets. Red and blue colors represent colon/rectum and ileum samples, respectively. Circles and triangles represent anti-TNFα responders (TNF-R) and anti-TNFα inadequate responders (TNF-IR), respectively.

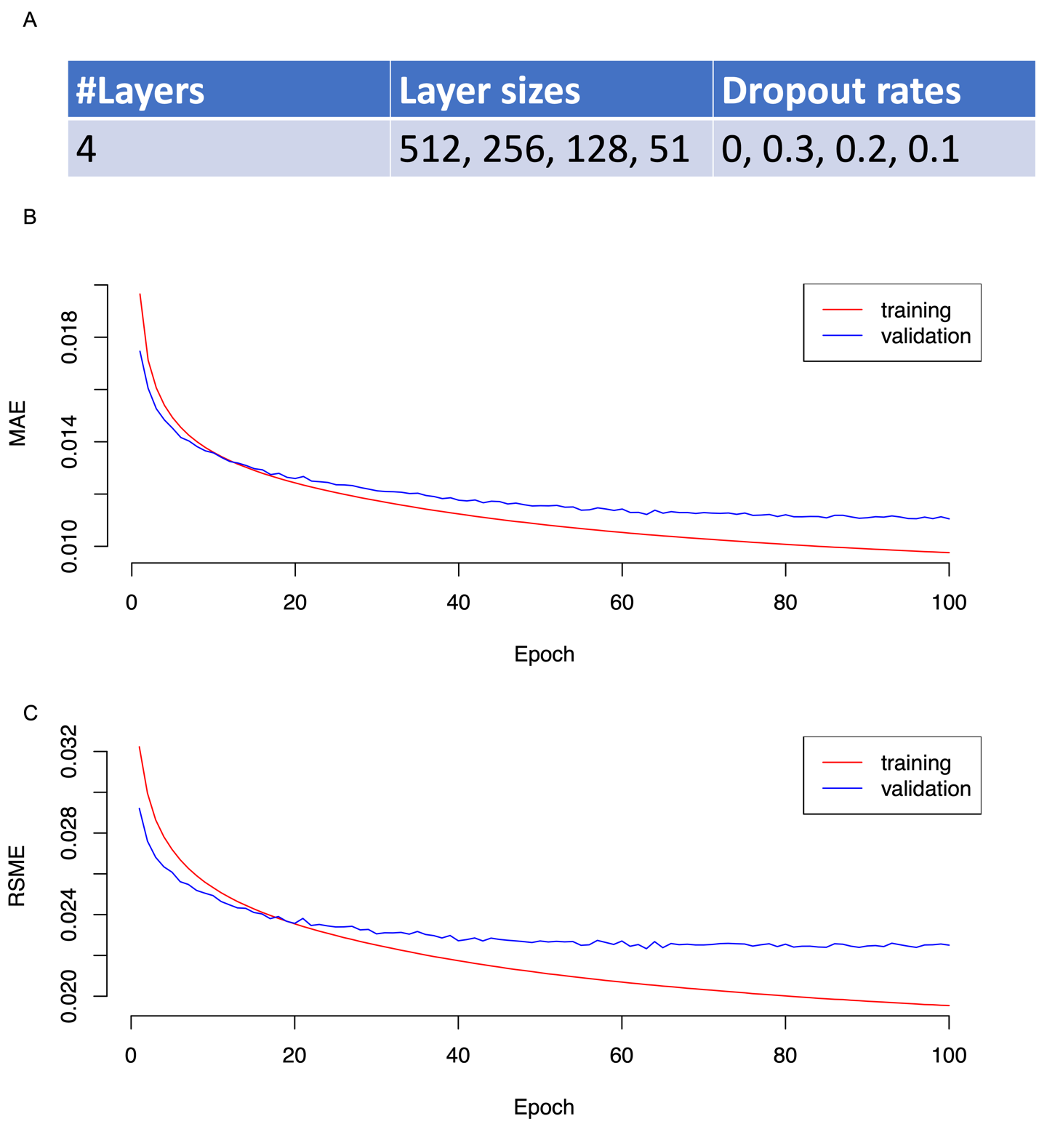

Supplementary Figure 2. Parameters (A) and performance (B-C) of deep-learning model trained by UC single-cell RNAseq data. Mean Absolute Error (MAE) and Root Mean Squared Error (RSME) were calculated based on the leave-one-subject cross-validation.

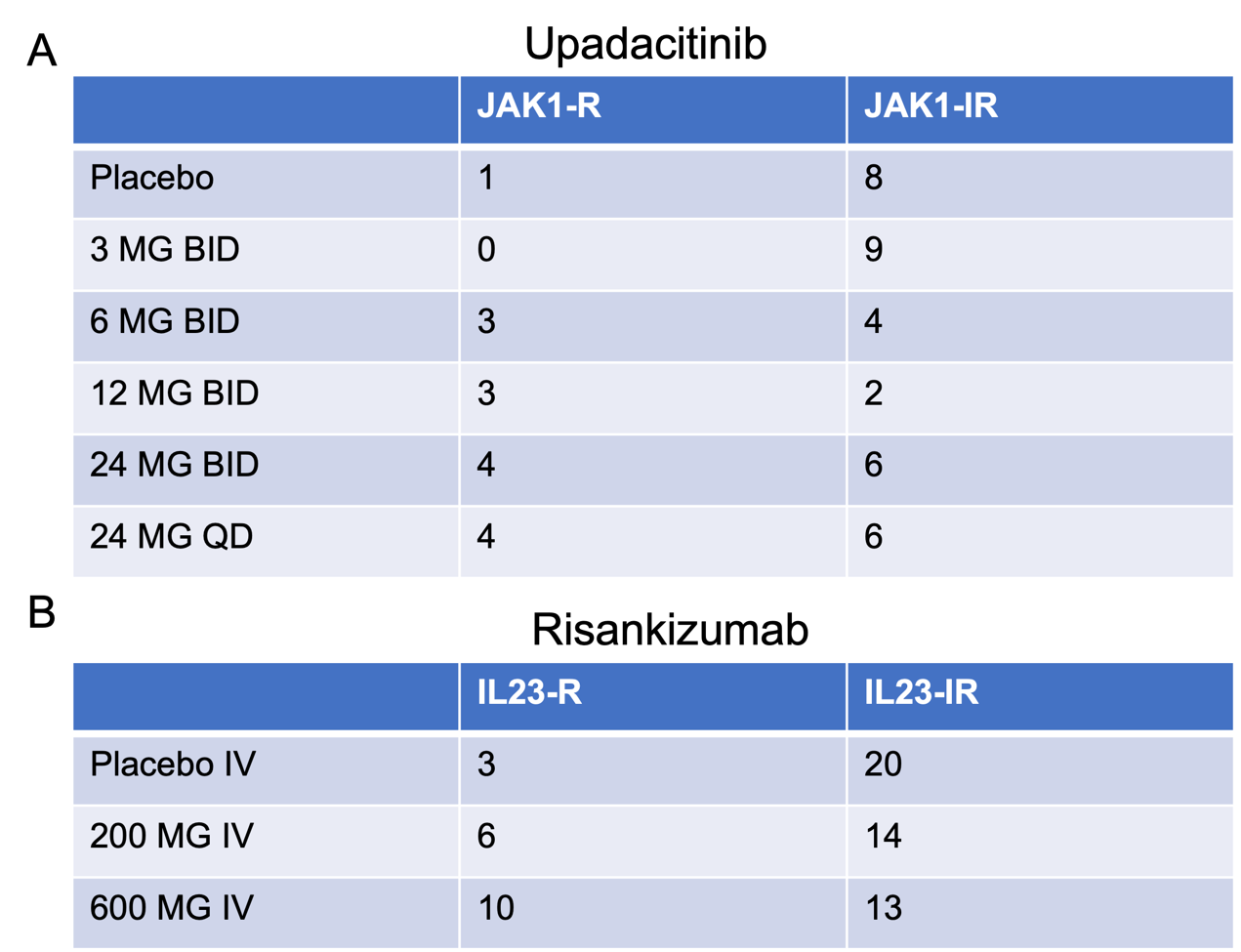

Supplementary Figure 3. Upadacitinib (A) and Risankizumab (B) clinical trial data used in our study. For upadacitinib, each number in the table represents the number of TNF-IR patients with paired week 0 and week 12/16 samples from colon biopsies in each arm. For risankizumab, each number in the table represents the number of TNF-IR patients with paired week 0 and week 12 samples from colon biopsies in each arm.

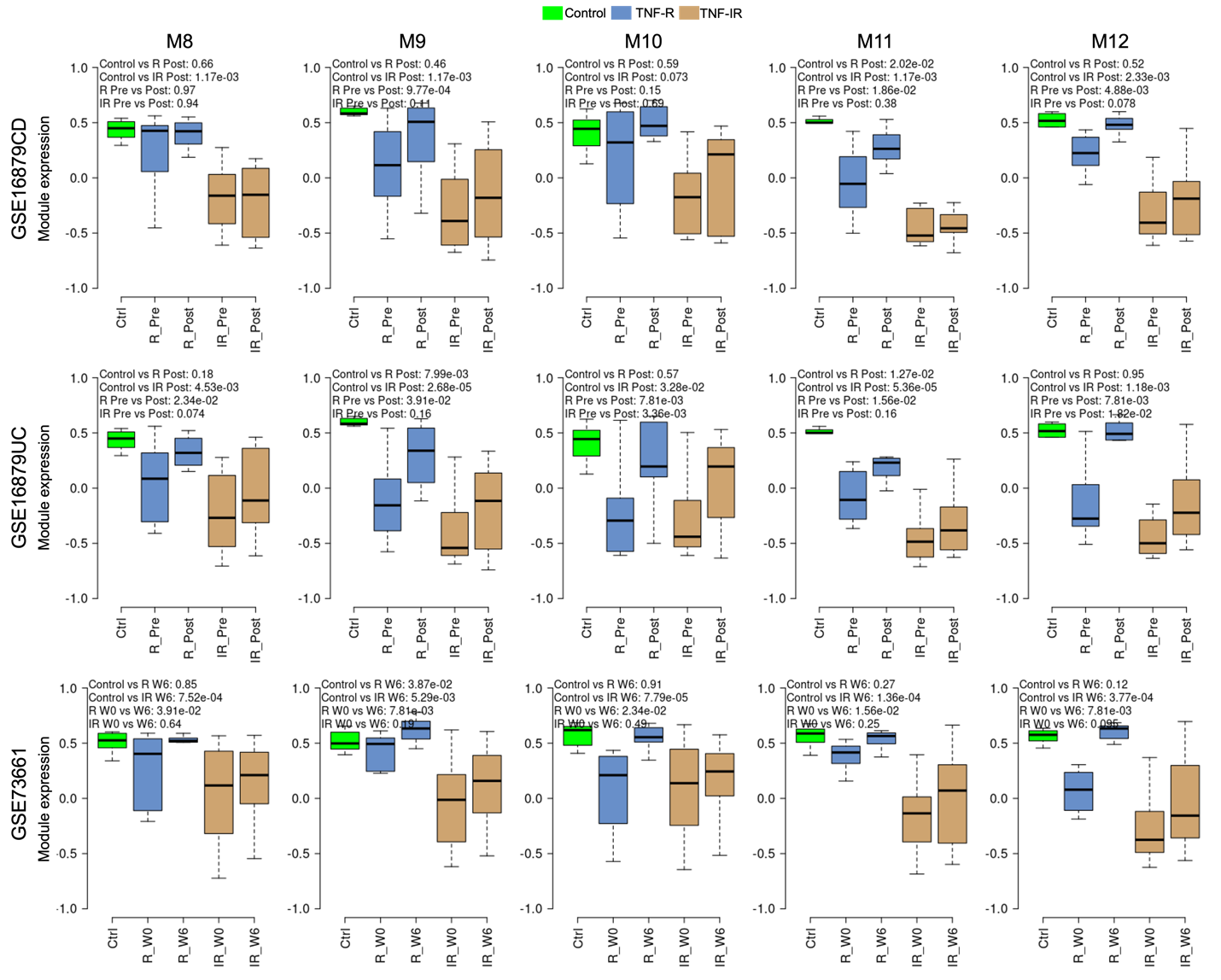

Supplementary Figure 4. The expression of TNF-IR down-regulated modules could not be returned to normal levels after anti-TNFα treatment in IR patients based on the GSE16879CD (A), GSE16879UC (B) and GSE73661 (C) datasets. Green, blue and light brown colors represent controls, TNF-R and TNF-IR patients, respectively. The p-values in the top-left of boxplots were calculated based on Wilcoxon rank sum test.

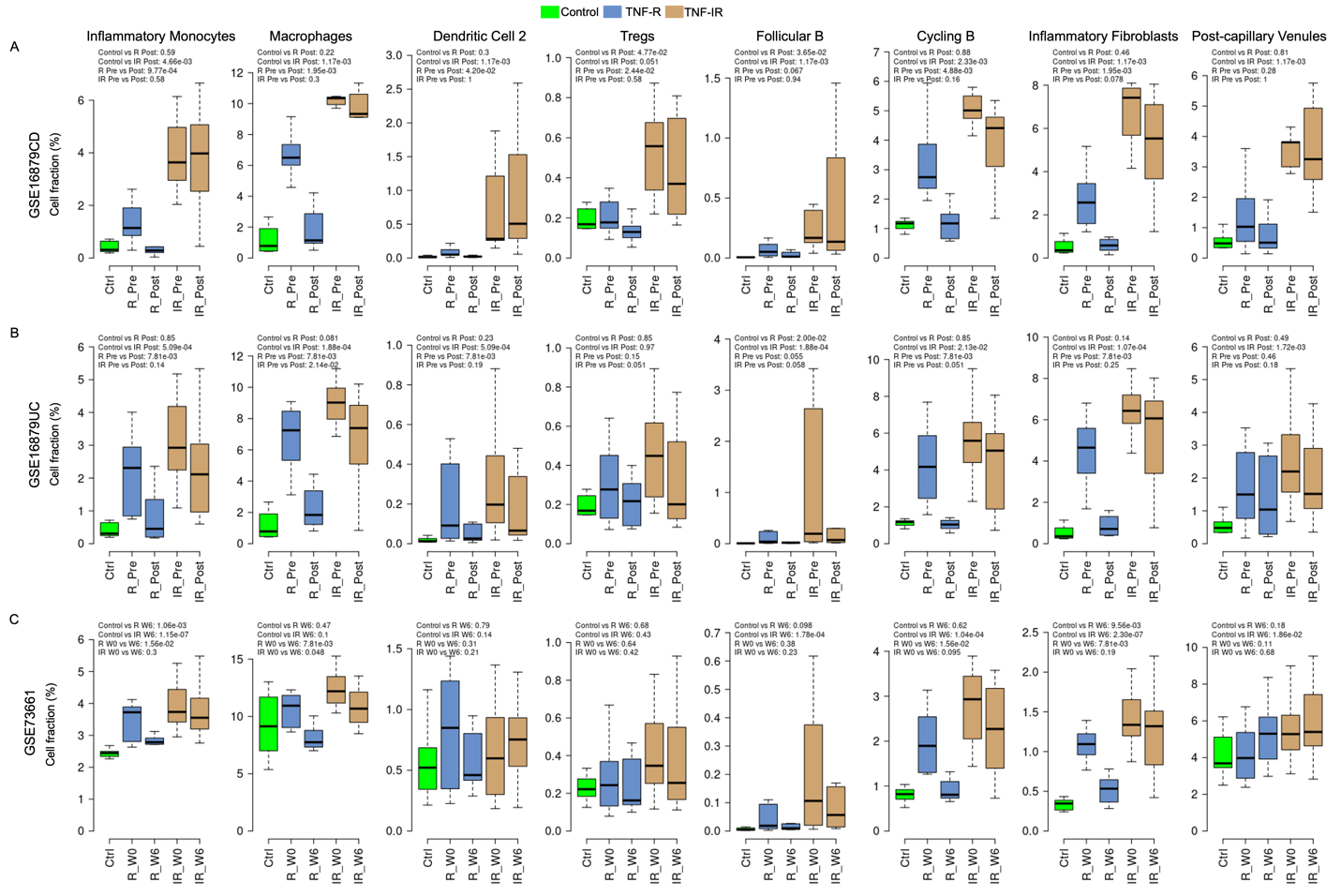

Supplementary Figure 5. Validation of TNF-IR up-regulated cell types. Green, blue and light brown colors represent controls, TNF-R and TNF-IR patients, respectively. The p-values in the top-left of boxplots were calculated based on Wilcoxon rank sum test.

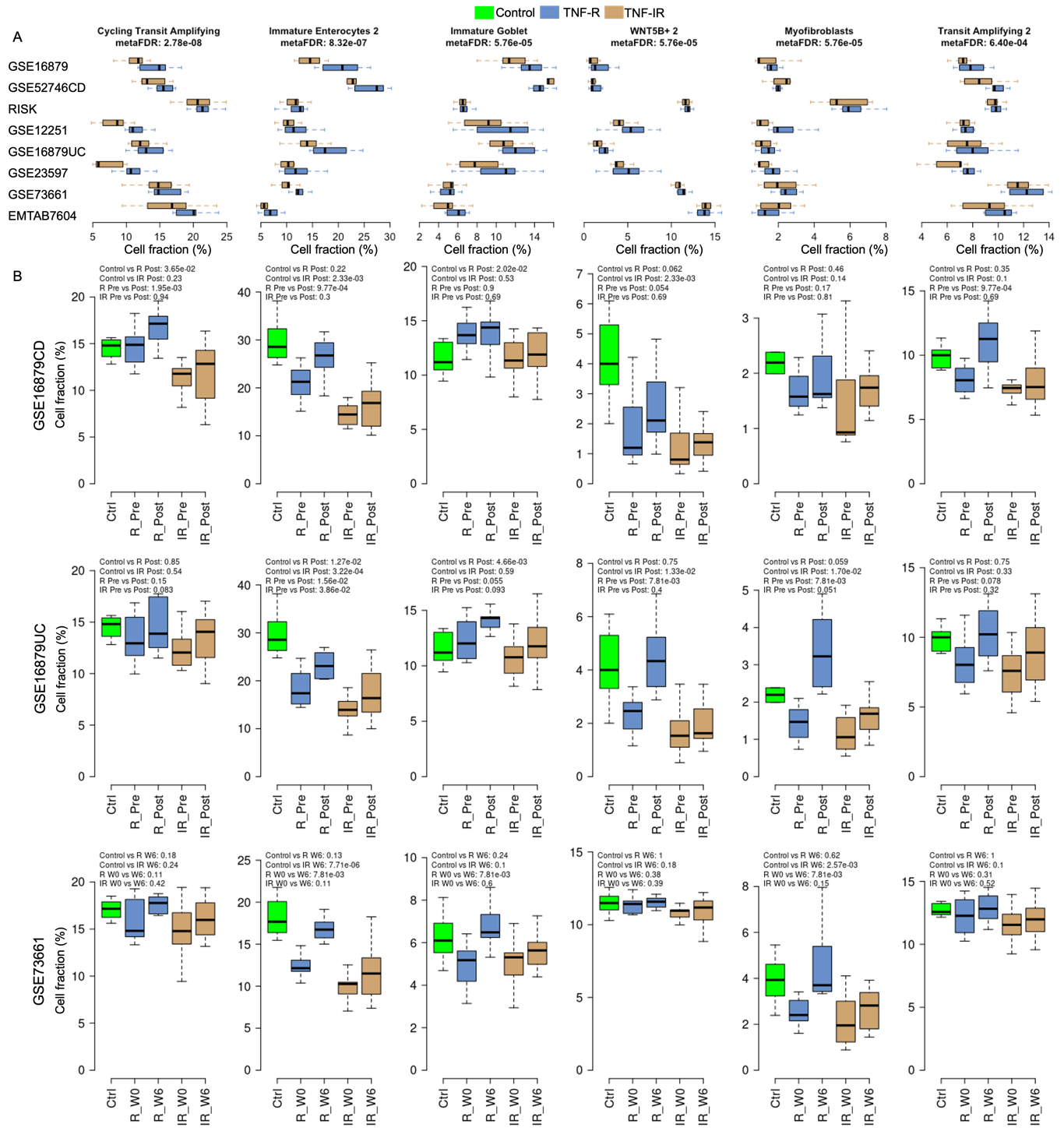

Supplementary Figure 6. Identification (A) and validation (B-D) of TNF-IR down-regulated cell types. Meta-FDR in (A) is calculated based on the comparison of cell fractions between TNF-IR and TNF-R patients in eight data sets. Green, blue and light brown colors represent controls, TNF-R and TNF-IR patients, respectively. The p-values in the top-left of boxplots in (B-D) were calculated based on Wilcoxon rank sum test.

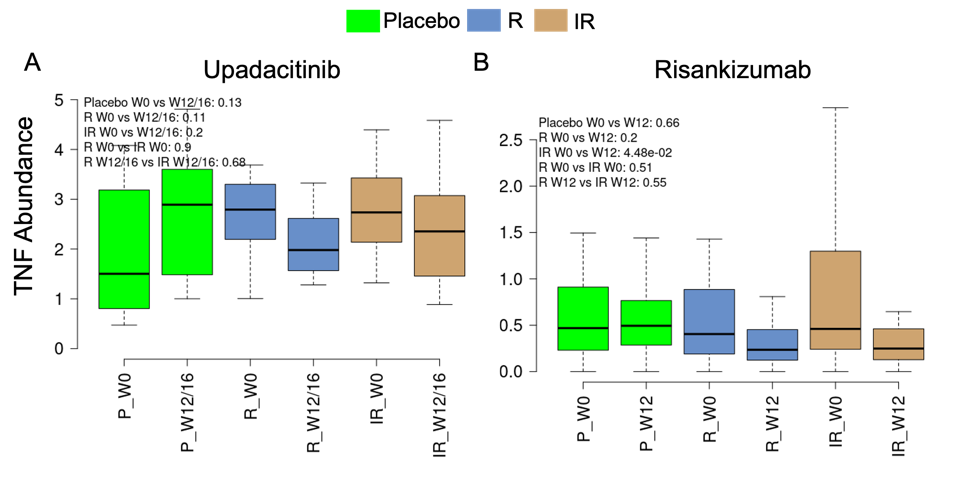

Supplementary Figure 7. Comparisons of TNF mRNA abundance between week 0 and week 12/16 for Upadacitinib among patients treated with placebo, JAK1-R patients and JAK1-IR patients (A) or between week 0 and week 12 for Risankizumab among patients treated with placebo, IL23-R patients and IL23-IR patients (B). Because over 80% of CPM (count per million) were less than 1 for TNF in Risankizumab RNAseq dataset, TNF was filtered out before performing the TMM normalization (see Supplementary Methods). Thus, CPM was used in (B). Green, blue and light brown colors represent placebos, JAK1-R (or IL23-R) and JAK1-IR (or IL23-IR) patients, respectively. The p-values were calculated based on paired or unpaired Wilcoxon rank sum test.

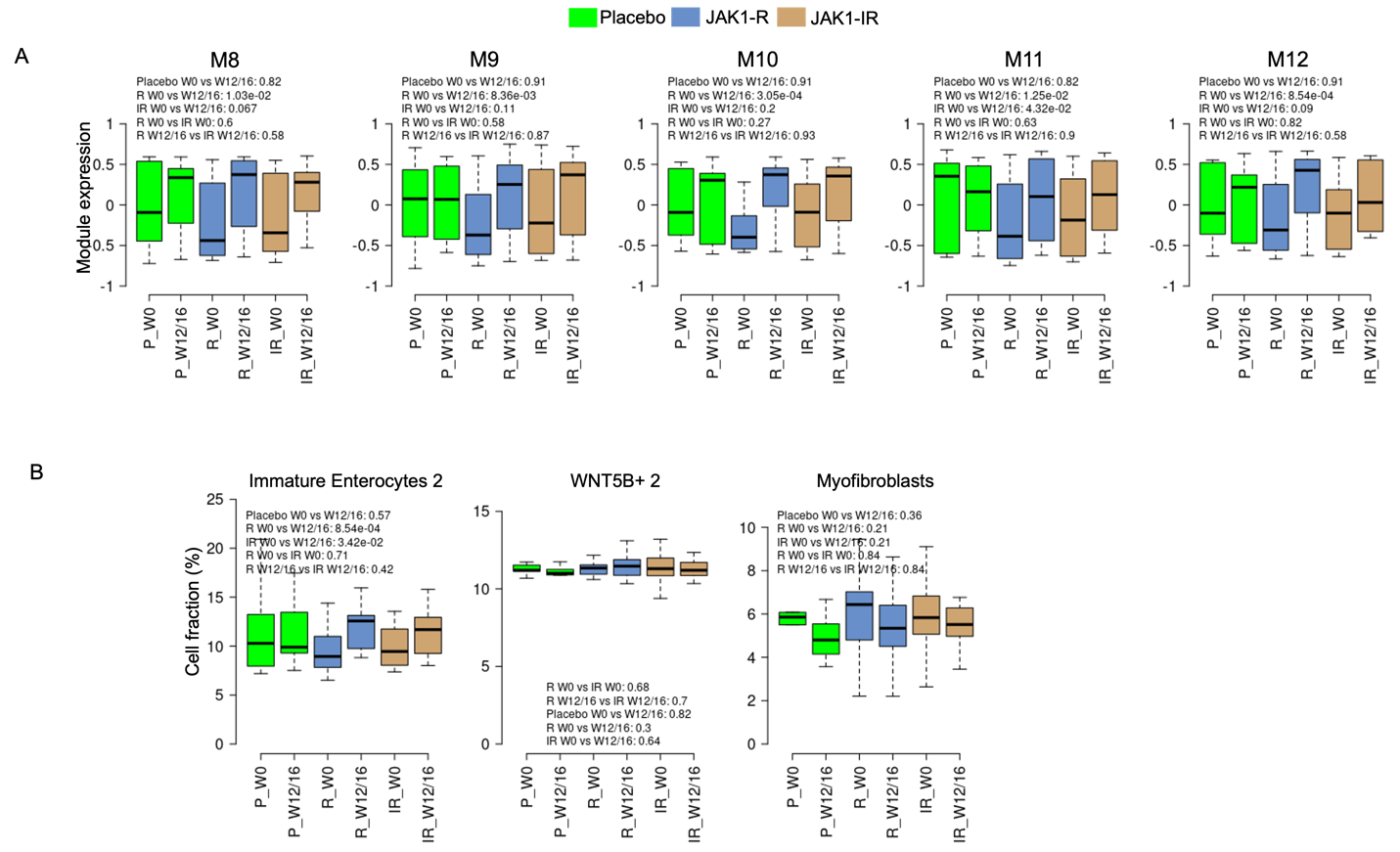

Supplementary Figure 8. Upadacitinib significantly affected TNF-IR down-regulated modules in JAK1-R patients but not placebo and JAK1-IR patients. (A) Comparison of expression of five TNF-IR down-regulated modules in placebos (green), JAK1-R (blue) and JAK1-IR (light brown) patients for week 0 and week 12/16. (B) Comparison of cell fractions of three TNF-IR down-regulated cell types in three groups for week 0 and week 12/16. The p-values were calculated based on Wilcoxon rank sum tests.

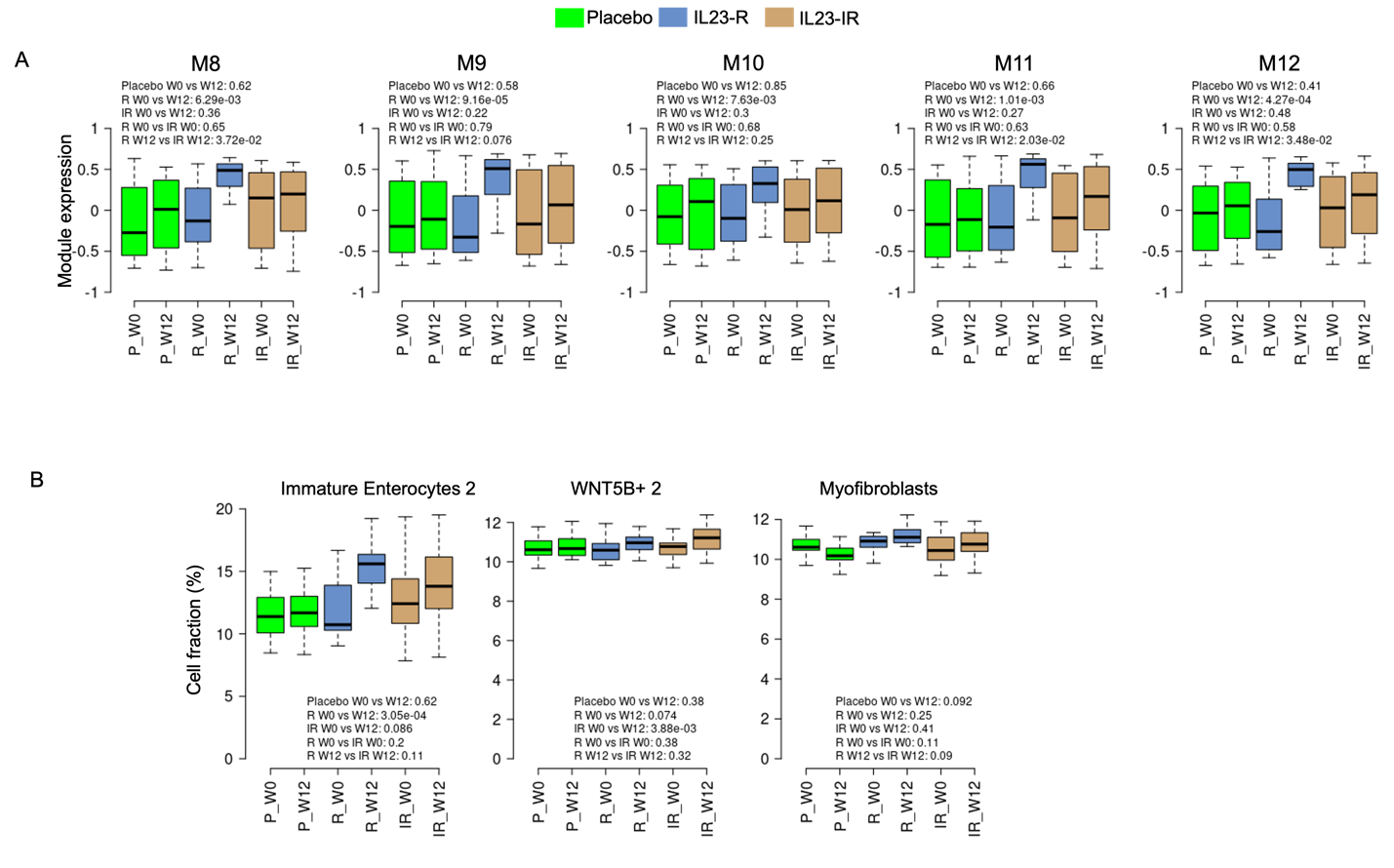

Supplementary Figure 9. Risankizumab significantly affected IL23-related TNF-IR down-regulated modules and cell types in IL23-R patients but not placebo and IL23-IR patients. (A) Comparison of expression of five TNF-IR down-regulated modules in placebos (green), IL23-R (blue) and IL23-IR (light brown) patients for week 0 and week 12. (B) Comparison of cell fractions of three TNF-IR down-regulated cell types in three groups for week 0 and week 12. The p-values were calculated based on Wilcoxon rank sum tests.

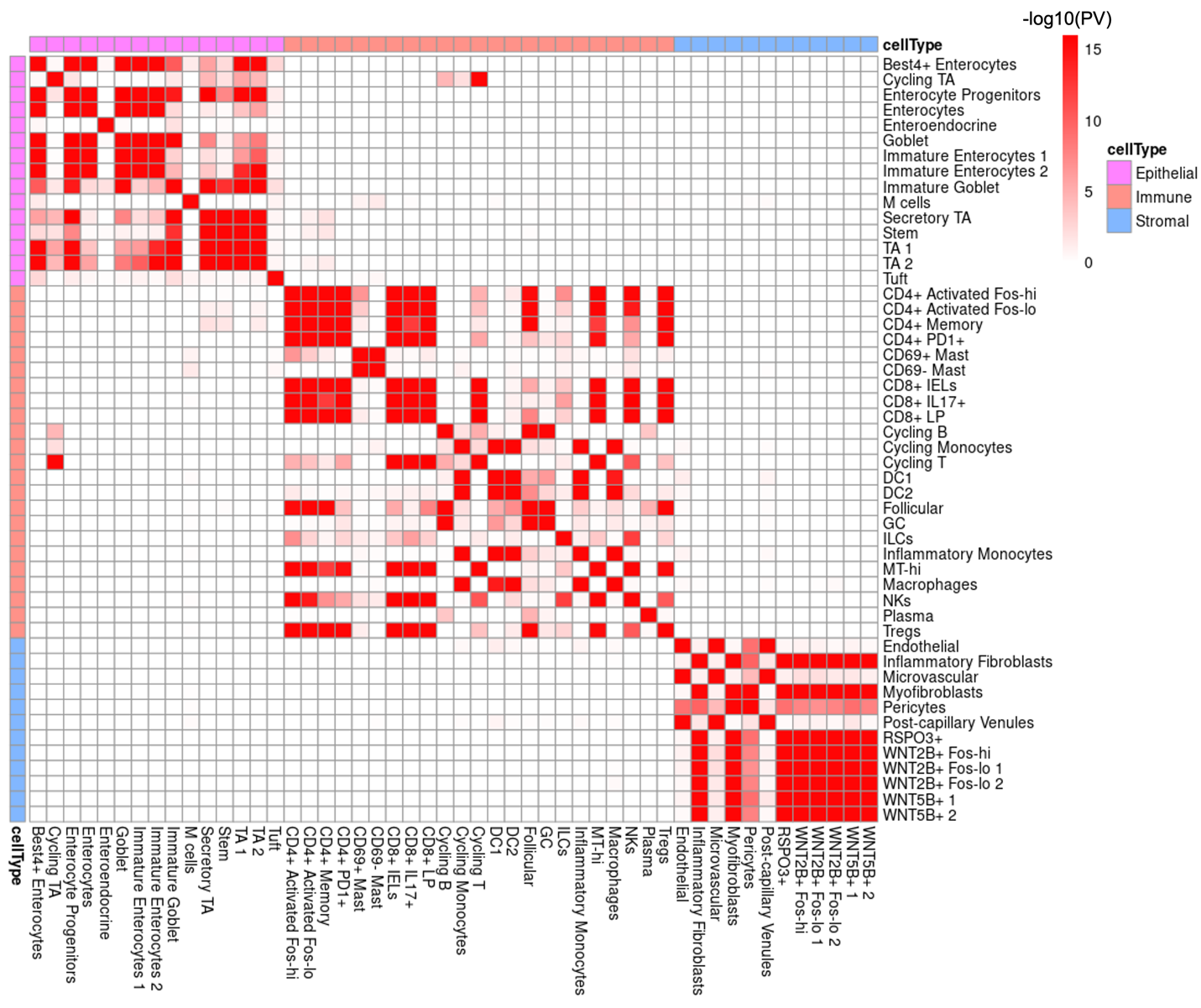

Supplementary Figure 10. Cell signature comparisons among cells from UC single-cell RNAseq data. The p value between each pair of cell signatures was calculated by hypergeometric test and then transformed by -log10. Red color in the heatmap represents the significant overlap between two cell signatures while white color represents no relationship between cell signatures.

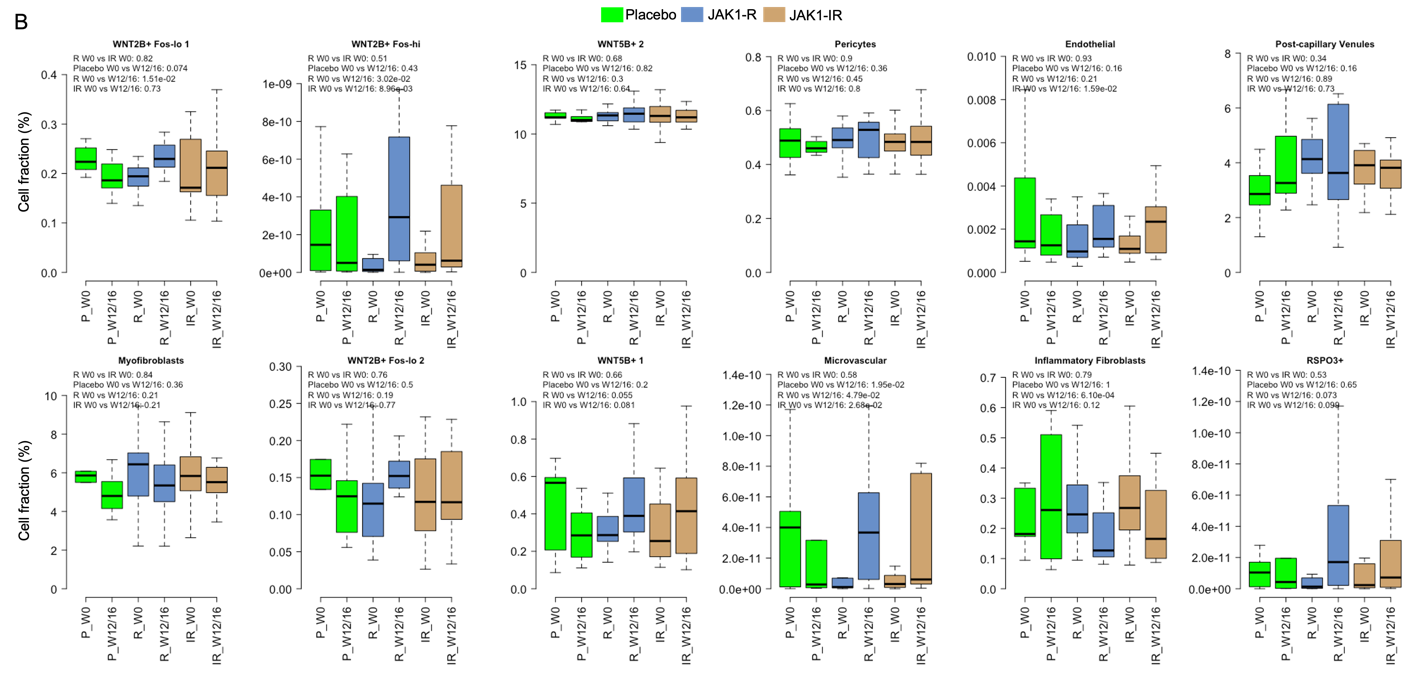

Supplementary Figure 11. Cell fraction comparisons of all stromal cells from UC single-cell RNAseq data between week 0 and week 12/16 samples for patients treated with placebos, JAK1-R patients and JAK1-IR patients. Green, blue and light brown colors represent placebos, JAK1-R and JAK1-IR patients, respectively. The p-values were calculated based on Wilcoxon rank sum tests.

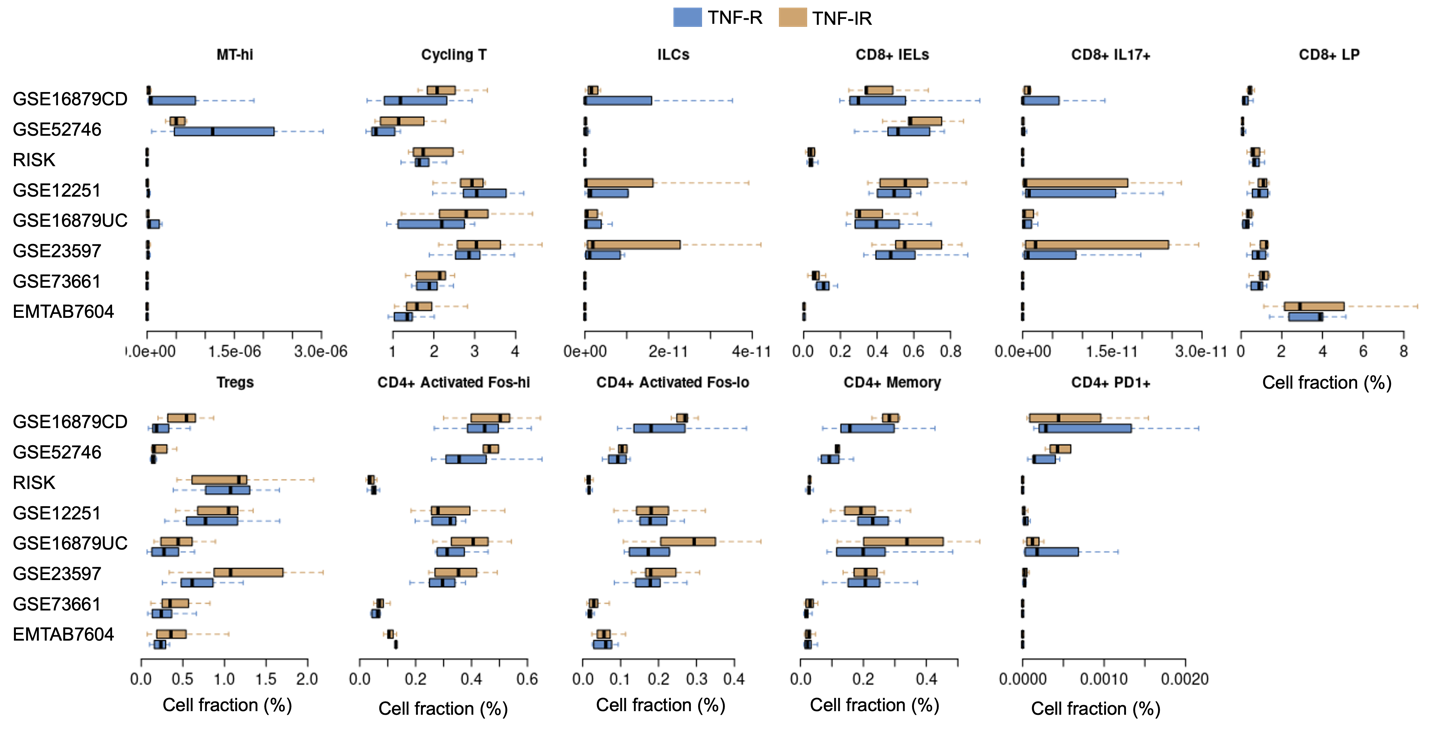

Supplementary Figure 12. Cell fractions comparisons of all T cells in UC single-cell RNAseq data between TNF-R and TNF-IR patients in eight datasets. Blue and light brown colors represent TNF-R and TNF-IR patients, respectively.

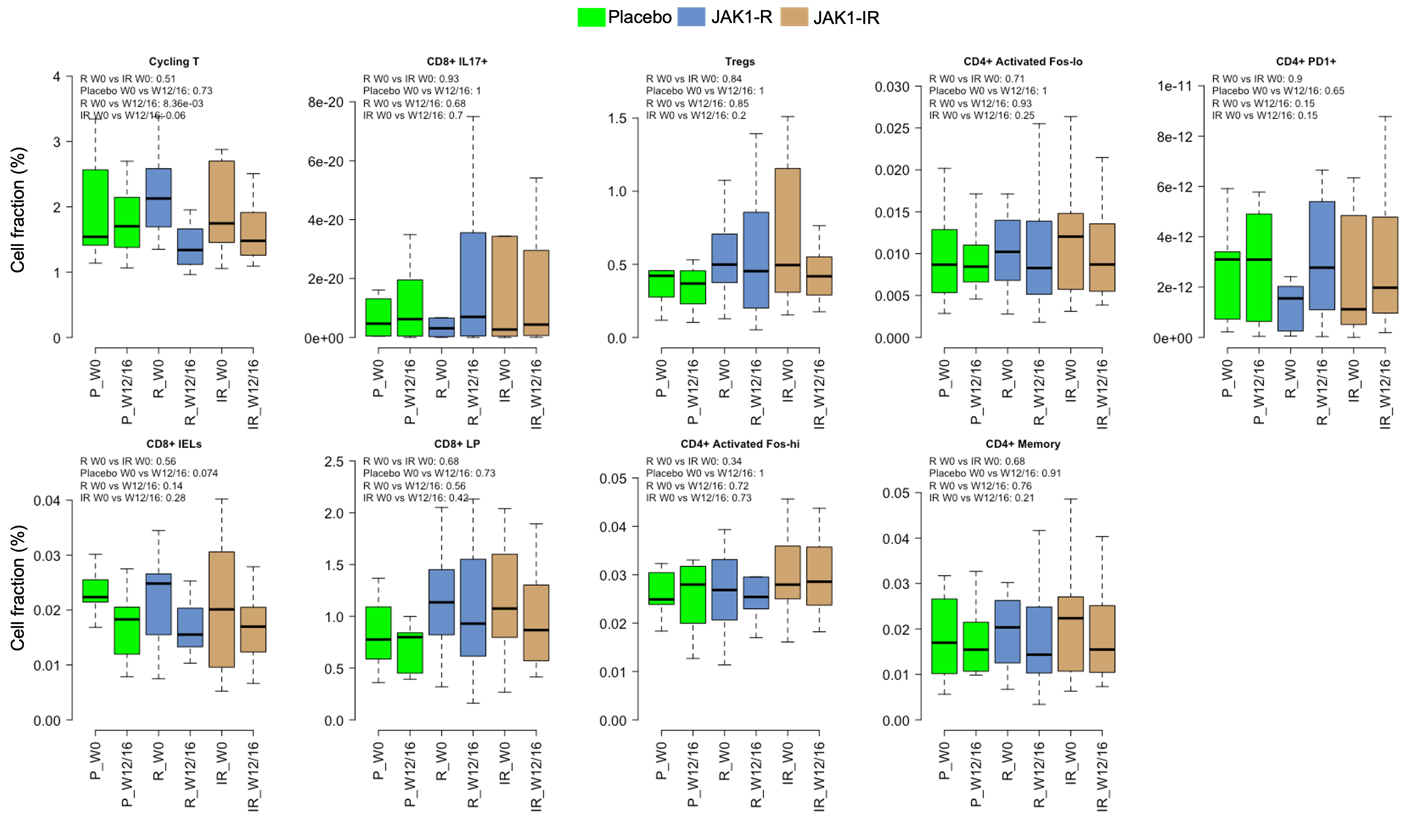

Supplementary Figure 13. Cell fraction comparisons of all T cells from UC single-cell RNAseq data between week 0 and week 12/16 samples for patients treated with placebos, JAK1-R patients and JAK1-IR patients. Green, blue and light brown colors represent placebos, JAK1-R and JAK1-IR patients, respectively. The p-values were calculated based on Wilcoxon rank sum tests.
