## Supplementary Methods for "The Clinical Response of Upadacitinib and Risankizumab is Associated with Reduced Inflammatory Bowel Disease Anti-TNFα Inadequate Response Mechanisms"

**Anti-TNFα gene expression data processing**

The Affymetrix raw data from GEO data sets were processed by the R package affy for RMA normalization and log transformation. Probe IDs were mapped to the HUGO gene symbols based on a mapping table from GEO. For RNA-seq data E-MTAB-7604 and RISK, quantifiable genes were filtered by CPM>1 in at least 20% of samples and then the count matrix was normalized and log transformed using the EdgeR package^1^.

**Meta-P and meta-FDR calculation**

Two-sided p-values were converted to two one-sided p-values (TNF-IR up-regulated p-value and down-regulated p-value) based on the directionality of log2 fold change.

$$Upregulated p=\left\{ \begin{matrix} \begin{matrix} p/2 & if log2\left( fold change \right)>0 \end{matrix} \\ \begin{matrix} 1-p/2 & otherwise \end{matrix} \end{matrix} \right.$$

$$Downregulated p=1-upregulated p$$

A meta p-value for the TNF-IR up-regulated p-values or down-regulated p-values of a gene in all data sets was calculated by the following equation from the R package “metap”:

$$metaP= \frac{\sum_{i=1}^{k} z(p_{i})}{\sqrt{k}}$$

where k was the number of TNF-IR up-regulated (or down-regulated) p-values (k≤8 because a gene may not be included in all data sets). A meta False Discovery Rate (FDR) was calculated by the BH (Benjamini-Hochberg) method from the p.adjust function in R.

One limitation of a meta p-value calculation occurs when a one-sided p-value from one data set is highly significant (e.g. 1E-10), although the directionality of the same gene in several other data sets was inversed, the meta p-value may also be significant. To avoid this limitation, consensus DEGs were identified based on two rules: TNF-IR up-regulated (or down-regulated) meta-FDR<0.05 and one-sided p-value<0.2 in at least 5 data sets.

**Consensus WGCNA modules identification**

Based on the consensus DEGs, scale-free topology fit index (R^^2^) and mean connectivity for each soft thresholding power were first calculated by the pickSoftThreshold function in R package WGCNA^2^. The minimum power was then selected under R^^2^≥0.8 and mean connectivity ≤ 30 based on the suggestion from Langfelder et al ^3^ (GSE12251 power was selected based on R^^2^=0.794 and mean connectivity=14.1). An adjacent matrix for each data set was then calculated based on the power-transformed correlation between each pair of genes. Because genes with a negative correlation did not correspond to the functional similarity^4^, we only included positive correlations in the adjacency matrix. Topology overlap matrix (TOM)^2^ was calculated based on the adjacent matrix. Because TOM matrices had different statistical properties, a quantile normalization method suggested by Langfelder et al.^5^ was used to scale the TOM matrices such that the 95^th^ percentile of all TOM matrices were equal.

$${scaledTOM}_{i}={{TOM}_{i}}^{{scaledQuan}_{i}} i\in[1,8]$$

$${scaledQuan}_{i}=\frac{log({Quantile95}_{1})}{log({Quantile95}_{i})} i\in[1,8]$$

where Quantile95_i_ was the 95% quantile of 200,000 values randomly selected from TOM_i_. A consensus TOM matrix was generated by calculating the mean of the scaled TOM score of each pair of genes in all scaled TOM matrices. Because consensus DEGs may not be included in all data sets, scaled TOM scores of missed gene pairs in each data set were assigned as 0 when generating the consensus TOM matrix. Finally, consensus modules were identified based on the consensus TOM matrix by WGCNA R package

**Deep-learning based cell fraction estimation based on single-cell RNAseq data**

Menden et al.^6^ developed Scaden (single cell-assisted deconvolutional deep neural network (DNN)), which can build a deep-learning model based on pseudo-bulk RNA-seq samples generated from single-cell RNAseq data and then predict cell type fractions for the input samples of cell mixtures. The advantages of Scaden compared with other single-cell based cell deconvolution methods (e.g. MUSIC^7^ or CIBERSORTx^8^) relate to the non-linear combined hidden features of DNN, which make results more robust for the input noise and technical bias. Thus, based on the method description and parameter suggestion from the original study, we developed the R version Scaden to deconvolute eight anti-TNFα treatment bulk data sets based on the UC single-cell RNASeq data generated by Smillie et al.^9^.

UC single-cell RNASeq data was downloaded from the Broad Institute single cell portal (<https://singlecell.broadinstitute.org/single_cell>) and included 51 cell clusters with a total of 295,442 cells from colon mucosa of 18 UC patients with adjacent inflamed and non-inflamed biopsies and 12 healthy controls. The single-cell data was processed by “Seurat” analysis pipeline^10^. Based on the recommendation of Menden et al.^6^, each pseudo-bulk sample only included cells from one single-cell sample, which captured cross-sample heterogeneity. 50,000 pseudo-bulk samples were generated from 47 single-cell samples in the UC single-cell RNAseq data (non-inflamed sample from patient N12 had less than 1,000 cells and was removed). The number of pseudo-bulk samples generated from each single-cell sample was based on the percentage of cells in this single-cell sample among cells in all single-cell samples.

The cell types with at least 30 cells in a single-cell sample were selected to generate pseudo-bulk samples and each pseudo-bulk sample included at least 5 cell types with a total of 500 cells from this sample. Further details about generating each pseudo-bulk sample can be found from Menden et al.^6^.

Based on 50,000 pseudo-bulk samples, we built the 4-layer DNN model with L1 as a loss of function and Rectified Linear Unit (ReLU) activation for all layers but the last and softmax activation for the last layer (Supplementary Figure 2A). The model was trained and validated based on the leave-one-out subject method developed by Menden et al.^6^. Because MAE (mean absolute error) and RSME (Root Mean Square Error) of the validation cohort was already less than 0.012 and 0.024 (Supplementary Figure 2B and 2C), we did not generate three DNN models with different parameters and calculate the averaged predictions from these models like Menden et al.^6^. To estimate the cell fractions of real bulk data, the real data were logarithmically transformed, and each sample was scaled to the range [0,1].
